# Microtubule networks in zebrafish hair cells facilitate presynapse transport and fusion during development

**DOI:** 10.1101/2024.04.12.589161

**Authors:** Saman Hussain, Katherine Pinter, Mara Uhl, Hiu-Tung Wong, Katie S. Kindt

**Affiliations:** Section on Sensory Cell Development and Function, National Institute on Deafness and other Communication Disorders, Bethesda, MD, 20892, USA; Presynaptogenesis and Intracellular Transport in Hair Cells Junior Research Group, Institute for Auditory Neuroscience and InnerEarLab, University Medical Center Goettingen, 37075 Goettingen, Germany; Collaborative Research Center 889 ‘Cellular Mechanisms of Sensory Processing’, 37075 Goettingen, Germany

## Abstract

Sensory cells in the retina and inner ear rely on specialized ribbon synapses for neurotransmission. Disruption of these synapses is linked to visual and auditory dysfunction, but it is unclear how these unique synapses form. Ribbon synapses are defined by a presynaptic density called a ribbon. Using live imaging in zebrafish hair cells, we find that numerous small ribbon precursors are present throughout the cell early in development. As development progresses, fewer large ribbons remain, and localize at the presynaptic active zone (AZ). Using tracking analyses, we show that ribbon precursors exhibit directed motion along an organized microtubule network to reach the presynaptic AZ. In addition, we show that ribbon precursors can fuse together on microtubules. Using pharmacology, we find that microtubule disruption interferes with ribbon motion, fusion, and normal synapse formation. Overall, this work demonstrates a dynamic series of events that underlies the formation of a critical synapse required for sensory function.

## Introduction

The inner ear and retina contain sensory cells with specialized ribbon synapses that faithfully transmit the timing, duration, and intensity of sensory stimuli to the brain. These synapses are critical for hearing, balance, and vision, and their disruption is linked to auditory, vestibular, and visual disorders (Frederick and Zenisek, 2023; Kujawa and Liberman, 2015; Wan et al., 2019).

The hallmark feature of these synapses is the presynaptic ribbon, a dense body made up primarily of the protein Ribeye (Schmitz et al., 2000). Ribbons are found at the presynaptic active zone (AZ) and act as scaffolds to ready synaptic vesicles for release (Schmitz, 2009). Work in fixed tissues has led to the hypothesis that small ribbon precursors migrate to the AZ and fuse to form larger, mature ribbons. However, direct evidence for these dynamic processes during ribbon formation has not yet been demonstrated.

Neurotransmission at mature ribbon synapses is triggered in response to graded membrane depolarizations dictated by the duration and intensity of sensory stimuli. Membrane depolarization opens voltage-gated calcium channels (Ca_V_1) beneath ribbons (Brandt et al., 2003; Chang et al., 2006) (see schematic in Figure 1C). Calcium influx triggers synaptic vesicle fusion and the release of glutamate onto postsynaptic receptors (Obholzer et al., 2008; Ruel et al., 2008). Studies have shown Ribeye, the core component of the ribbon, is essential for the formation and function of ribbon synapses in both mouse and zebrafish (Becker et al., 2018; Jean et al., 2018; Lv et al., 2016; Maxeiner et al., 2016). Other key components include the classic neuronal scaffolding proteins Bassoon and a novel variant of Piccolo, Piccolino (Gundelfinger et al., 2016). In mice, the loss of Bassoon disrupts ribbons anchoring and synapse function (Dick et al., 2003; Frank et al., 2010; Khimich et al., 2005). In rats, the loss of Piccolino also impacts ribbon morphology (Michanski et al., 2023; Regus-Leidig et al., 2014). Currently how these key molecular players fit within the dynamics underlying ribbon formation is unclear.

**Figure 1.**
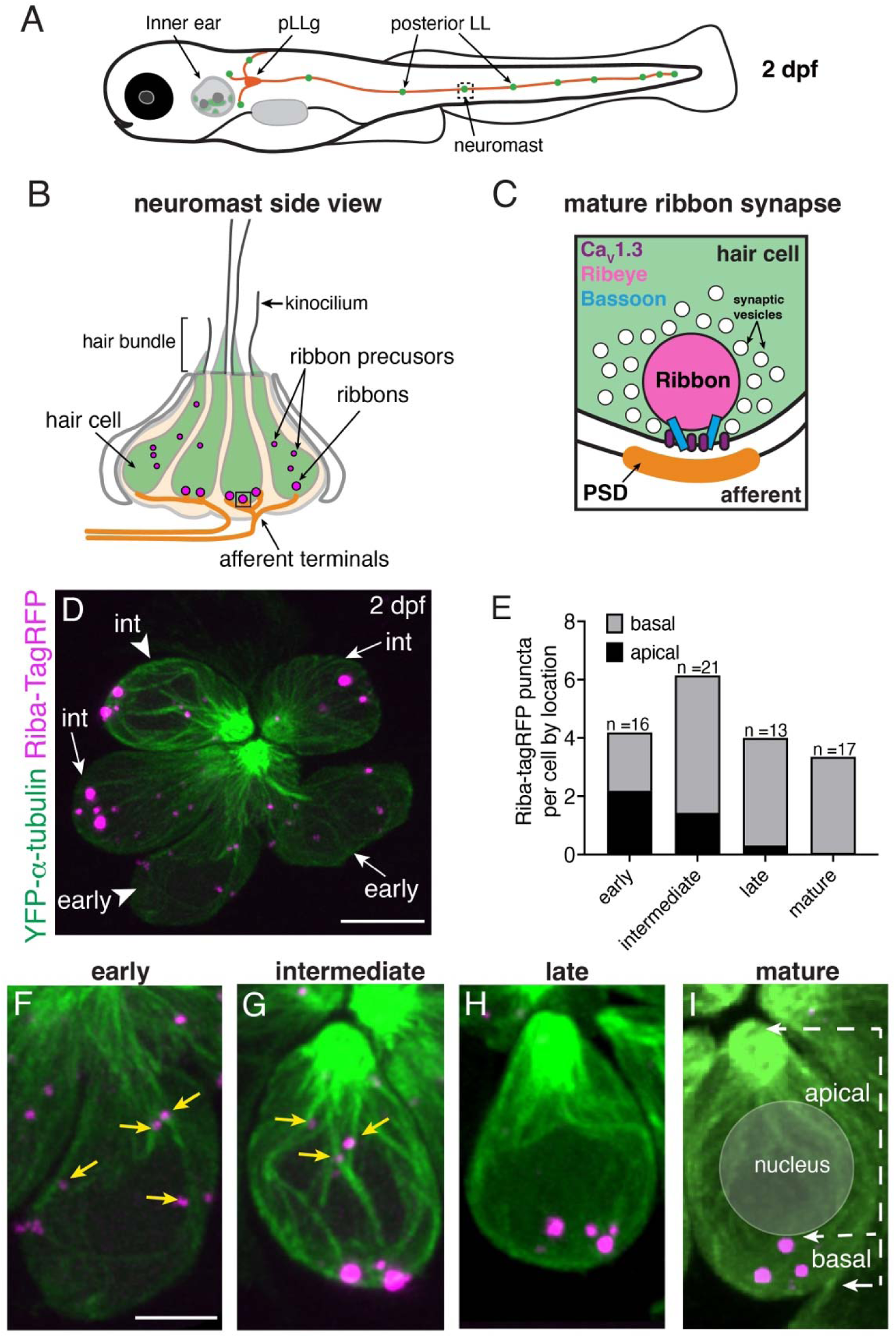
Ribbons associate with microtubules and change localization during development. **A)** Schematic of a larval zebrafish at 2 days post fertilization (dpf) with the location of the posterior-lateral line (pLL) indicated relative to the inner ear. Neuromasts (green) in the posterior LL contain sensory hair cells that are innervated by afferent projections from the posterior LL ganglion (pLLg, orange). **B)** Schematic of a neuromast at 2 dpf, viewed from the side. At 2 dpf, the majority of hair cells (green) are developing. At the top of the cells, the mechanosensory hair bundle is composed of actin-based stereocilia and a tubulin-based kinocilium. A shorter kinocilium and an abundance of small ribbon precursors are indicative of an immature stage. **C)** Schematic of a ribbon synapse when mature. The dense presynapse or ribbon is made primarily of Ribeye. Ribbons tether synaptic vesicles near Ca_V_1.3 channels at the plasma membrane, across from the postsynaptic density (PSD). Bassoon acts to anchor ribbons at the presynaptic AZ. **D)** Example image of a neuromast at 2 dpf, viewed from top down. The microtubule network and ribbons are marked with YFP-Tubulin and Riba-TagRFP respectively. In this example of 6 developing hair cells, 2 early and 4 intermediate cells are present. The cell bodies of an early and intermediate cell from this example (arrowheads) are expanded in **F** and **G**. **E)** Plot shows the average number of ribbons per hair cell at each developmental stage. Cell stage is determined by the height of the kinocilium. After an increase in ribbon number with development, there is a decrease upon maturation. The number of apically-localized Riba-TagRFP puncta is high at early and intermediate stages and is lower in late and mature hair cells. In contrast, the number of basally-localized Riba-TagRFP puncta is low at early stages and becomes higher by intermediate stages (n = 16, 21, 13, 17 hair cells for early, intermediate, late, and mature stages respectively). **F-I)** Example images of hair cells expressing YFP-Tubulin and Riba-TagRFP at early, intermediate, late, and mature stages. At the early stage, Riba-TagRFP puncta are spread throughout the cell body and are smaller in size. At the intermediate stage, the number of Riba-TagRFP puncta becomes more basally enriched and are larger in size. In late and mature hair cells, all Riba-TagRFP puncta are at the base of the cell and are fewer in number compared to the intermediate stage. The arrows in **I** highlight the apical and basal region of the cell used for quantification of Riba-TagRFP puncta location in **E**. Yellow arrows in **E** and **F** indicate precursors associated with microtubules. Scale bars in **D** = 5 µm and in **F** = 2 µm.

In neurons, precursor vesicles containing Piccolo and Bassoon are transported along microtubules to the developing presynaptic AZ. (Ahmari et al., 2000; Gundelfinger et al., 2016; Maas et al., 2012; Shapira et al., 2003). This transport requires molecular motors, along with adaptor proteins (Bury and Sabo, 2011; Fejtova et al., 2009; Maas et al., 2012). In neurons, kinesins are the molecular motors that transport cargo in the anterograde direction–towards the presynaptic AZ–while cytoplasmic dynein mediates retrograde transport, back to the cell soma (Sweeney and Holzbaur, 2018). Although the kinesin responsible for Piccolo-Bassoon vesicle transport is not known, recent work in *Drosophila* and *C. elegans* suggests that the kinesin KIF1A may transport AZ components (Oliver et al., 2022; Pack-Chung et al., 2007). Work in developing photoreceptors and mouse auditory inner hair cells (IHCs), has shown that ribbon precursors contain not only Ribeye, but also Bassoon, and Piccolino (Regus-Leidig et al., 2009; Michanski et al., 2019). Whether ribbon precursors are actively transported along microtubules during synapse formation is not known.

The formation of ribbon synapses has been extensively studied using light and electron microscopy in fixed tissues (Michanski et al., 2019; Regus-Leidig et al., 2009; Schmitz, 2009; Sheets et al., 2011; Sobkowicz et al., 1986, 1982). In mouse auditory IHCs, this process occurs over an extended time period (E18-P14) (Michanski et al., 2019; Sobkowicz et al., 1986, 1982), while in zebrafish hair cells, ribbon synapses mature in just 12-18 hrs (Dow et al., 2015; Sheets et al., 2011). Early development in both mouse IHCs and zebrafish hair cells features many small ribbon precursors throughout the cell, likely formed via Ribeye self-aggregation in the cytosol (Magupalli et al., 2008; Schmitz et al., 2000). As development progresses, ribbons enlarge, localize to the presynaptic AZ, and associate with the innervating afferent terminals. Finally, the number of ribbons associated with postsynaptic machinery is refined to obtain the proper number of complete synapses. Recent work in mice has shown that ribbon precursors associate with microtubules (Michanski et al., 2019), and it has been proposed that ribbon precursors may migrate along microtubules to reach the presynaptic AZ, although the *in vivo* dynamics of this process remain unclear.

To study ribbon formation, we examined hair cells and developing ribbons in the zebrafish lateral line (Figure 1A), a sensory system that allows aquatic vertebrates to sense local water movements (Freeman, 1928; Suli et al., 2012). The lateral line consists of clusters of hair cells called neuromasts that are arranged in lines along the surface of the fish. In the posterior lateral line (pLL), which forms an array of neuromasts along the zebrafish trunk, hair cells emerge at 2 days post fertilization (dpf) (see Figure 1A-B,D), and the lateral-line system reaches functional maturity by 5 dpf (Suli et al., 2012). At these larval ages zebrafish are transparent, which enables *in vivo* imaging of ribbon formation over extended periods (Dow et al., 2015). Further, transgenic lines expressing fluorescently tagged proteins allow high-resolution visualization of subcellular dynamics, including ribbon formation.

During our study, we worked in close collaboration with another group investigating the late stages of ribbon formation in mouse auditory IHCs (postnatally). This work is published in a companion paper (Voorn et al., 2024). Together our studies demonstrate that ribbon transport along microtubule networks is essential for proper ribbon formation in mice and zebrafish. Our zebrafish work leverages transgenic lines that label: developing ribbons, microtubule networks, and the growing plus ends of microtubules. We use these lines, along with high-resolution imaging to visualize the dynamics of ribbon formation. Using live imaging, we show that early in development, many small ribbon precursors are distributed throughout the cell. By later stages, fewer large ribbons remain and localize to the base of the cell. We show that microtubule networks in lateral-line hair cells are dynamic and grow plus ends towards the presynaptic AZ, the preferred direction for most kinesin motors (Sweeney and Holzbaur, 2018). Tracking analyses reveal the directed motion of ribbon precursors towards the presynaptic AZ, with ribbon precursors moving along and fusing on microtubules. Further, an intact microtubule network is critical for ribbon transport, fusion, and synapse formation. Overall, this foundational work provides insight into ribbon formation and the processes needed to reform ribbon synapses for the treatment of auditory and vestibular synaptopathies.

## Results

### Time course of ribbon formation in living zebrafish lateral-line hair cells

Fixed preparations in zebrafish and mice have outlined a conserved process that underlies ribbon or presynapse development in hair cells (Michanski et al., 2019; Sheets et al., 2011). To examine the time course underlying ribbon formation *in vivo*, we studied hair cells in the zebrafish pLL. These hair cells form 3-4 ribbon synapses in just 12-18 hrs (Dow et al., 2015; Kindt et al., 2012). To image hair cells, ribbon precursors, and ribbons *in vivo* we used a double transgenic line. One transgenic line labels microtubules and serves as a marker of hair cells (*myo6b:YFP-tubulin*). The other transgenic line reliably labels ribbons and smaller ribbon precursors (*myo6b:riba-TagRFP*) in hair cells (see example: Figure 1D). Although this latter transgene expresses Riba-TagRFP under a non-endogenous promoter, neither the tag nor the promoter ultimately impacts cell numbers, synapse counts, or ribbon size (Figure 1-S1A-E).

For our initial analyses, we assessed the overall time course of ribbon formation in hair cells in the posterior lateral line (pLL) when larvae were 2 dpf (Figure 1A). At 2 dpf each pLL neuromast contains 4-8 developing hair cells at different developmental stages (Figure 1B,D,F-I). We staged hair cells based on the development of the apical, mechanosensory hair bundle. The hair bundle is composed of actin-based stereocilia and a tubulin-based kinocilium. We used the height of the kinocilium (see schematic in Figure 1B), the tallest part of the hair bundle, to estimate the developmental stage of hair cells as described previously (stage: hair bundle height; early: < 1.5 µm, intermediate: 1.5-10 μm, late: 10-18 μm, mature: > 18 µm (Zhang and Kindt, 2022)). Qualitatively, we observed that at early and intermediate stages, small Riba-TagRFP puncta were present throughout the cell body (see examples: Figure 1D,F-G). By late stages, or in newly matured hair cells, larger Riba-TagRFP puncta were restricted to the base of the cell (see example: Figure 1H-I). We quantified the total number of Riba-TagRFP puncta based on developmental stage and found that the total number of puncta was high at early and intermediate stages but significantly decreased when hair cells were mature (Figure 1E, Figure 1-S2A, n = 7 neuromasts and 67 hair cells).

When taking these live images, we had no postsynaptic marker and were unable to differentiate between precursors and smaller ribbons associated with a postsynaptic density. Therefore, we classified all apical Riba-TagRFP puncta above the nucleus as precursors, and all ribbons located beneath the nucleus at the cell base as more mature ribbons. Using this classification, we quantified the total number of Riba-TagRFP puncta at each developmental stage (Figure 1E, Figure 1-S2B-C, n = 7 neuromasts and 67 hair cells). We found that apical Riba-TagRFP precursors were only present at early and intermediate stages (Figure 1E, Figure 1-S2B, and see yellow arrows in Figure 1F-G). In contrast, hair cells at late or mature stages contained more mature ribbons located at the cell base, below the nucleus (Figure 1E,H-I, Figure 1-S2C). Overall, our live images examining Riba-TagRFP puncta show a similar, developmental process as observed in fixed preparations–the number of ribbons and precursors decrease and become basally localized to the presynaptic AZ as the hair cell develops.

### Microtubules grow plus ends toward the presynaptic active zone in hair cells

An important question in ribbon formation is how ribbons and precursors migrate to the presynaptic AZ. Work on mouse IHCs using electron microscopy (EM) and super-resolution microscopy found that developing ribbons and precursors often associate with microtubules (Michanski et al., 2019). Similar to what was observed in mice using EM, in living, pLL hair cells we observed ribbon precursors associated with microtubules (see yellow arrows in Figure 1F-G). Based on these association studies, a microtubule network may function to transport ribbon precursors during development. To understand if ribbons could be transported along microtubules, we first examined the composition and polarity of the microtubule network in pLL hair cells.

We first examined the composition, or posttranslational modifications present in the microtubule network in pLL hair cells using immunohistochemistry. Using this approach, we labeled either acetylated-(modification found in mechanically stabilized microtubules) or tyrosinated-(modification found in destabilized microtubules) α-tubulin (Janke and Magiera, 2020). We performed our staining in a transgenic line that labels all microtubules (*myo6b:YFP-tubulin*). We found that in pLL hair cells, the microtubule network extends along the apical-basal axis of the cell and is highly acetylated; acetylated microtubules are more concentrated at the cell apex (Figure 2-S1A-C). Outside of the kinocilium, we did not observe tyrosinated microtubules in pLL hair cells. Instead, tyrosinated microtubules were observed in the zebrafish skin, pLL nerve terminals, and the supporting cells that surround hair cells (Figure 2-S1D-F).

Overall, our immunostaining results indicate that in pLL hair cells, a considerable portion of the microtubule network is highly acetylated. Acetylation may provide a population of mechanically stable microtubules that could be used to transport ribbon precursors.

Although microtubule modifications are informative, these labels do not provide definitive information regarding microtubule growth or polarity. Knowing microtubule polarity is important, as many cargos are transported based on polarity. For example, most kinesin motor proteins transport cargo toward the more dynamic plus end of microtubules (Sweeney and Holzbaur, 2018). Therefore, to explore microtubule polarity in pLL hair cells, we created a transgenic line that expresses the plus-end marker of microtubules, EB3 (end-binding protein 3), fused with GFP (see example: Figure 2A, *myo6b:EB3-GFP*). Previous work in other cell types has shown that EB3-GFP can be used to visualize the plus end of growing microtubules (Kawano et al., 2022; Stepanova et al., 2003).

**Figure 2.**
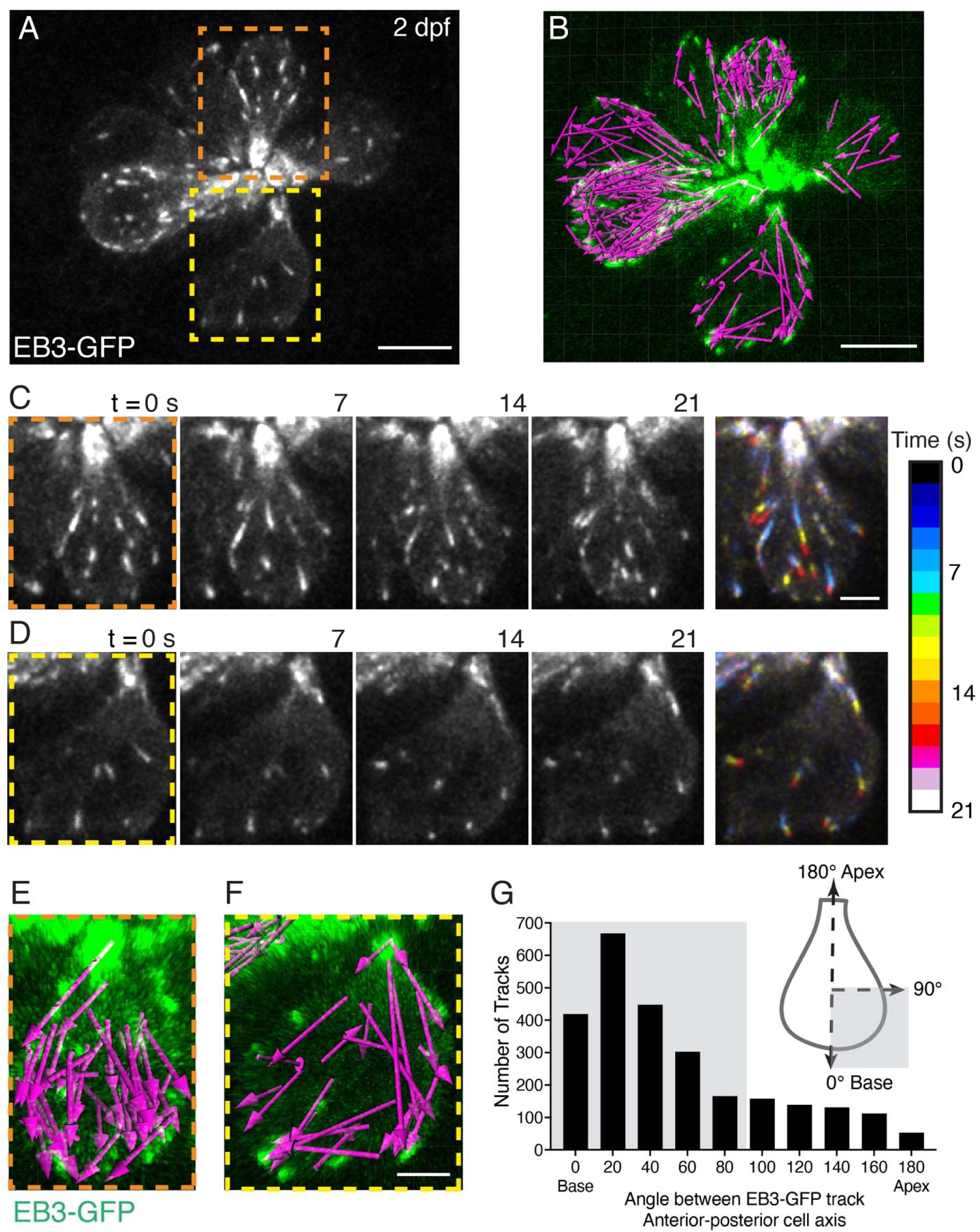
EB3-GFP tracks show plus ends of microtubules move to the cell base. **A** Example image of a neuromast at 2 dpf. The growing or plus ends of microtubules are marked with EB3-GFP. In this example, the apex of 8 developing cells is at the center of the image and the base of each cell is at the periphery. 2 example cells are outlined (dashed lines) and expanded in more detail in in **C-D,** and **E-F**. **B)** A 22-min timelapse was taken of the example in **A**. All EB3-tracks, indicated by magenta arrows (tracked in Imaris) detected during the timelapse are shown. **C-D)** Example time courses of EB3-GFP in hair cells over 21 s; the cell apex is towards the top of each image. In the final image, the 4 images for each example (0-21 s) were projected over time as a pseudocolor image represented by the colormap. The pseudocolor images show that many EB3-GFP tracks move to the cell base. **E-F)** The magenta arrows in **E** and **F** show all the EB3-GFP tracks acquired in the example cells in **C-D** over the entire 22-min duration. Arrowheads indicate the direction of travel. **G)** The schematic in **G** shows how EB3-GFP tracks were aligned to each hair cell. Tracks moving toward the apex have a track angle of 180°, while those moving to the base have an angle of 0°. This analysis revealed that the majority of EB3-GFP tracks (shaded domains) move toward the base of the cell (n = 7 neuromasts, 2597 tracks). Scale bars in **A-B** = 5 µm and in **C-F** = 2 µm.

We used this transgenic line, along with confocal microscopy to visualize microtubule growth in pLL hair cells (z-stacks every 7 s for 20-30 min). By imaging EB3-GFP dynamics, we observed comet-like tracks that allowed us to visualize the plus end of growing microtubules (see Movie S1). In hair cells, microtubule organizing centers are located beneath a single microtubule-based kinocilium, the primary cilium in hair cells (Lepelletier et al., 2013).

Consistent with studies in motile cilia, we observed foci of EB3-GFP at the tips of kinocilia that emanate away from the apex of the cell body (Schrøder et al., 2011) (Figure 2-S2A-C). This result suggests that similar to other cilia, the plus ends of kinocilia are towards their tips, away from the cell body. In addition, we observed EB3-GFP tracks within the soma of hair cells and found that the majority of tracks were directed away from the cell apex and towards the base of the cell (see EB3-GFP images taken from Movie S1, with tracks rendered into a pseudocolor image based on time, Figure 2C-D). To quantify the direction of EB3-GFP tracks we used Imaris to perform 2D analyses to detect and create vectors of EB3-GFP tracks (see arrows; Figure 2B,E,F). We aligned these vectors along the 2D apical-basal axis of each cell. Here vector movement towards the apex was represented as 180°, and movement to the base was represented at 0° (see schematic, Figure 2G). The movement of EB3-GFP vectors was quantified as the angle between 0 and 180°. From this analysis, we found that 78 % of EB3-GFP vectors were directed towards the base in pLL hair cells (Figure 2G, 2014/2597 tracks < 90°, n = 7 neuromasts and 33 hair cells).

Overall, our immunostaining revealed that the soma of hair cells contains a population of microtubules stabilized by acetylation. Our *in vivo* imaging of EB3-GFP revealed that within this population there is extensive microtubule dynamics, and the plus ends of microtubules are primarily directed towards the cell base, and the presynaptic AZ.

### Ribbon precursors associate with and show directed motion along microtubules

Our analysis of microtubule dynamics using EB3-GFP indicates that the plus ends of microtubules point toward the cell base (Figure 2). Therefore, we hypothesized that these tracks of microtubules could be used by kinesin motor proteins to transport precursors from the cell apex to the base during development. To test this hypothesis, we used either Airyscan or Airyscan 2 confocal imaging to capture timelapses of ribbon and precursor movement for longer durations (Airyscan: ∼3 µm z-stacks (15-20 slices) every 50-100 s for 30-70 min) or for shorter total durations with a faster capture rate (Airyscan 2: ∼2-3.5 µm z-stacks (12-20 slices) every 3-20 s for 5-40 min). We focused our analysis on developing pLL hair cells at early and intermediate stages when ribbon precursors are abundant throughout the cell (Figure 1D,F-G). For this work we used Riba-TagRFP to mark ribbons and precursors, and YFP-tubulin to label microtubules and provide cellular context.

From our timelapses, we observed that similar to our initial live analyses (Figure 1F-G), ribbon precursors associate with microtubules and are not found free-floating and untethered in the cell (Movie S2, S3). We observed that precursors exhibited 3 main movement behaviors related to microtubule association. First, we observed that the majority of precursors associated with microtubules remained stationary or confined (see examples: Figure 3C top two panels, Figure 3D,E asterisks, Movie S4). Second, we also observed rapid movement of precursors–during this movement, precursors appeared to be in close association with microtubules (see examples: Figure 3C bottom 4 panels, Figure 3D,E yellow arrowheads, Figure 3-S1A,B, Movie S4, S5, S6). This movement along microtubules occurred bidirectionally, towards the cell apex and towards the base (Figure 3C, middle panels (arrows to base), bottom panels (arrows to apex), Figure 3D-E, Figure 3-S1A and Movie S4, S5 (to base), Figure 3-S1B and Movie S6 (to apex)). Third, during the timelapses, we often observed that precursors switched associations between neighboring microtubules (Figure 3-S1C-D, Movie S7, 2.8 switching events per neuromast, n = 10 neuromasts).

**Figure 3.**
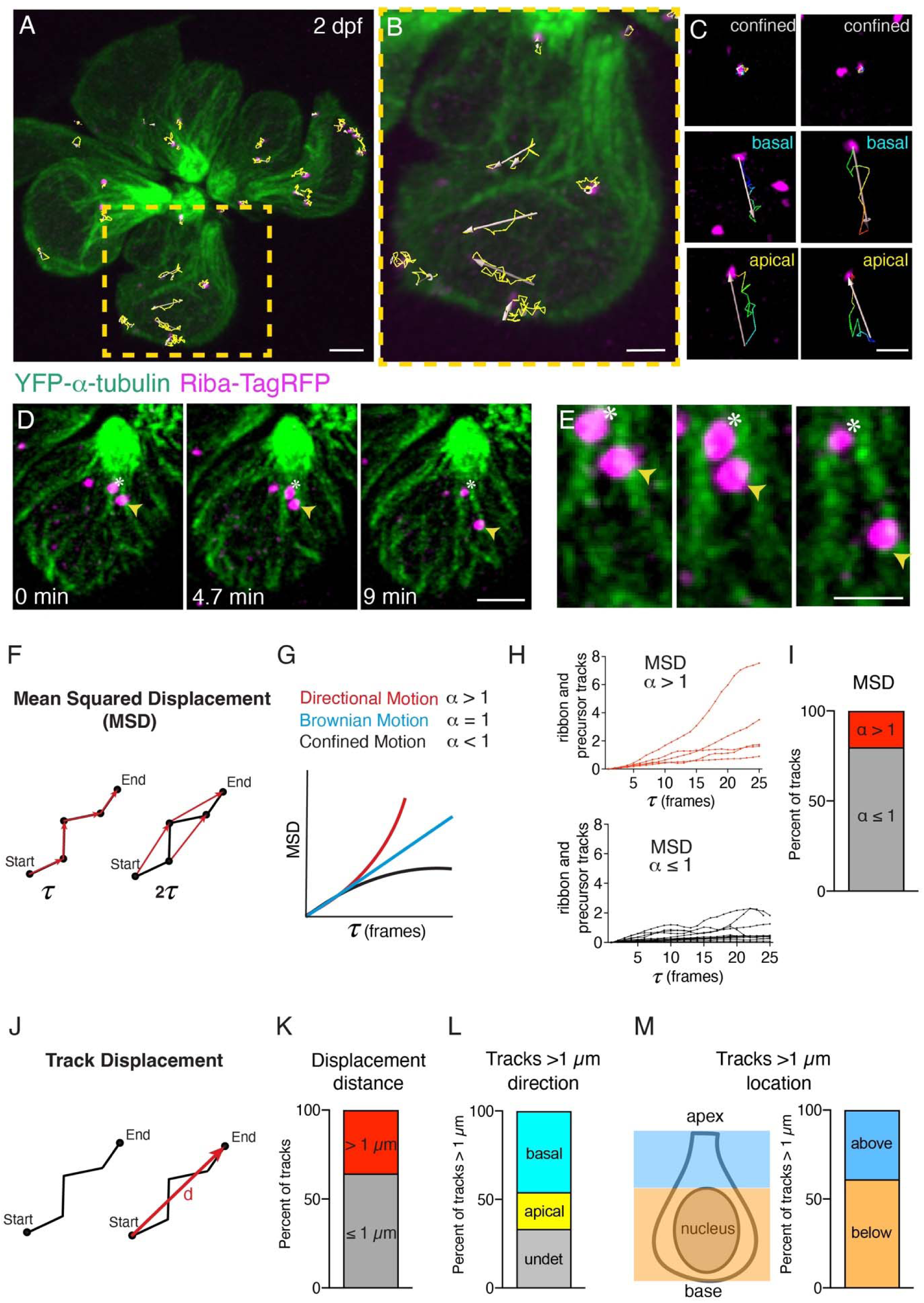
Ribbon precursors exhibit directional motion and confinement on microtubules. **A-B)** To quantify motion, ribbons and ribbon precursors were tracked in hair cells at early and intermediate stages at 2 dpf (example). Shown are tracks (in yellow) from an entire neuromast (**A**) and a single hair cell (**B**), obtained using Imaris, during a 30-min timelapse acquired every 50 s (also see Movie S2). **C)** Magnified view shows individual tracks over time and examples of confinement and motion towards the cell base and apex. **D)** Example of confined motion and directed motion on microtubules in a single hair cell. Movie was obtained during a 9 min timelapse acquired every 20 s. The ribbon labeled by the asterisks remains confined, while the ribbon labeled with the yellow arrow moves along a microtubule, towards the cell base, over time (also see Movie S4). **E)** A magnified image of the example shown in **D. F)** Mean squared displacement (MSD) vs time step was used to measure movement behaviors. Shown in red are the first- and second-time steps. The results are plotted in the form of MSD vs time step (**τ**) plots and the exponent (α) of the plots can be used to distinguish between the different types of motion observed (confined (α < 1), directional (α > 1), or Brownian motion (α = 1)). **H)** Example MSD plots of individual ribbon tracks from 2 control neuromasts (15 tracks MSD < 1 (black), 5 tracks MSD > 1 (red)). **I)** The bar graph shows the percent of MSD tracks displaying confined (α < 1, 79.8 %, gray), and directional motion (α > 1, 20.2 %, red). **J)** Track displacement vs time was used to measure movement behaviors with track > 1 µm indicative of directed motion. **K)** The bar graph shows the percent of tracks with distances > 1 µm (35.6 % red) and those with distances ≤ 1 µm (65.4 %, gray). **L)** The bar graph shows the percent of tracks with distances > 1 µm based on track direction (20.8 % to the cell apex, yellow; 45.8 % to the cell base, cyan; 33.3 % undetermined direction, gray). **M)** The bar graph shows the percent of tracks with distances > 1 µm based on location in the cell, above or below the nucleus (38.9 % above the nucleus, blue; 61.1 % below the nucleus, orange). In **K-M** n = 10 neuromasts, 40 hair cells, and 203 tracks. Scale bar in **A,D** = 2 µm and **B,C,E** = 1 µm.

To quantify ribbon and precursor movement, we used Imaris to obtain x,y,z coordinates for each ribbon during our timelapses (longer 30-70 min acquisitions) (see examples: Figure 3A-C, Movie S2, S3). This analysis yielded tracks or trajectories for all precursors and ribbons. We then performed a mean-squared displacement (MSD) analysis on ribbon tracks to classify the type of motion we observed. In this analysis, the exponent (α) of MSD vs time is obtained by curve fitting. A value of α > 1 indicates directional motion with velocity, α = 1 indicates Brownian motion, and α < 1 is representative of confined motion or subdiffusion (Figure 3F,G) (Sikora et al., 2017). This method allows us to determine what type of motion each track is exhibiting. Our results show that in developing hair cells, ribbon tracks exhibit directional as well as confined motion (Figure 3H, example MSD tracks with α > 1 (red tracks, top panel) and α < 1 (black tracks, bottom panel) from 2 neuromasts). Upon quantification, 20.2 % of ribbon tracks show α > 1, indicative of directional motion, but the majority of ribbon tracks (79.8 %) show α < 1, indicating confinement on microtubules (Figure 3I, n = 10 neuromasts, 40 hair cells, and 203 tracks).

To provide a more comprehensive analysis of precursor movement, we also examined displacement distance (Figure 3J). Here, as an additional measure of directed motion, we calculated the percent of tracks with a cumulative displacement > 1 µm. We found 35.6 % of tracks had a displacement > 1 µm (Figure 3K; n = 10 neuromasts, 40 hair cells, and 203 tracks). Of the tracks with displacement > 1 µm, the majority of ribbon tracks (45.8 %) moved to the cell base, but we also found a subset of ribbon tracks (20.8 %) that moved apically (33.4 % moved in an undetermined direction) (Figure 3L). This apical movement is consistent with a subpopulation of microtubules showing plus-end mediated growth apically (22.8 % of EB3-GFP tracks). In addition, we examined the location of precursors within the cell that exhibited displacements > 1 µm. We found that 38.9 % of these tracks were located above the nucleus, while 61.1 % were located below the nucleus (Figure 3M). Overall, our timelapse imaging demonstrated that ribbons and precursors displayed 3 main types of movement on microtubules: confinement, movement along microtubules, and switching between microtubules. Further, our tracking analyses indicate that while the majority of precursors are confined on microtubules, a subpopulation of ribbons and precursors exhibit directional motion.

### Long-term manipulation of microtubules impacts ribbon formation

Our timelapse imaging revealed that precursors and ribbons can move along directionally microtubules. To assess the importance of the microtubule network in synapse formation we used pharmacology to destabilize or stabilize the microtubule network in pLL hair cells using nocodazole or taxol respectively. We incubated larvae at 2 dpf for 16 hrs (56-72 hours post fertilization (hpf)), a time window that encompasses a large portion of synapse formation in developing hair cells. After these pharmacological treatments, we fixed and immunostained larvae to label acetylated-α-tubulin, to monitor changes to the microtubule network. In addition, we co-labeled with Ribeyeb to label ribbons and precursors and pan-Maguk to label postsynapses.

After a 16-hr treatment with 250 nM nocodazole, we observed a decrease in acetylated-α-tubulin label (qualitative examples: Figure 4A,C, Figure 4-S1A-B). Quantification revealed significantly less mean acetylated-α-tubulin label in hair cells after nocodazole treatment (Figure 4-S1D). Less acetylated-α-tubulin label indicates that our nocodazole treatment successfully destabilized microtubules. We also examined the number of hair cells per neuromast and observed that after nocodazole treatment, there were significantly fewer hair cells compared to controls. This indicates that either nocodazole is slightly toxic or interferes with cell division. Both situations have been observed previously during nocodazole treatments in other systems (Gupta, 1985; Zieve et al., 1980). We next examined the Ribeyeb label in hair cells to assess precursors and ribbons. We found that after nocodazole treatment, the total number of Ribeyeb puncta (apical and basal) per hair cell was significantly higher compared to controls (Figure 4G). We also observed that the average of individual Ribeyeb puncta (from 2D max-projected images) was significantly reduced compared to controls (Figure 4H). Further, the relative frequency of individual Ribeyeb puncta with smaller areas was higher in nocodazole-treated hair cells compared to controls (Figure 4I). We also examined the number of complete synapses (Ribeyeb-Maguk paired puncta) per hair cell after nocodazole treatment. We found that there were significantly fewer complete synapses per hair cell after a 16-hr nocodazole treatment (Figure 4F). Our long-term nocodazole treatment indicates that microtubule destabilization led to an increase in ribbons and precursors while reducing the number of synapses per cell.

**Figure 4.**
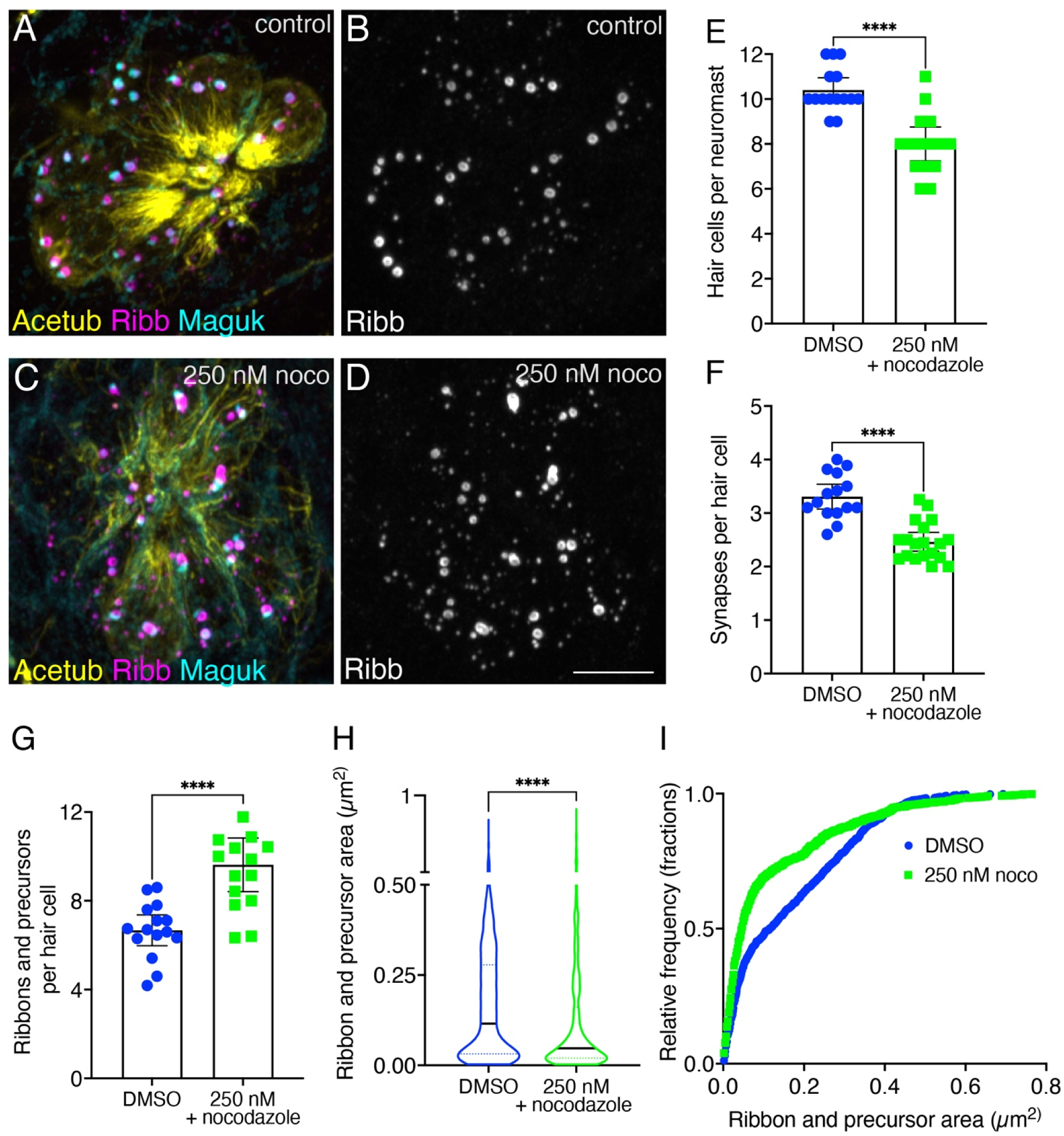
Overnight microtubule destabilization increases ribbon numbers, and decreases synapse counts and ribbon areas. **A-D)** Example immunostain of a neuromast at 3 dpf after an overnight treatment with 250 nM nocodazole (**C-D**) or DMSO (**A-B**). Acetylated-α-tubulin (Acetub) labels microtubules, Ribeyeb (Ribb) labels precursors and ribbons, and Maguk labels postsynapses. **E-G**) After an overnight treatment with 250 nM nocodazole there are significantly fewer hair cells per neuromast (**E**, P < 0.0001), fewer complete synapses per cell (**F**, P < 0.0001), and more ribbons and precursors per cell (**G**, P < 0.0001) compared to controls (n = 15 neuromasts for control and 250 nM nocodazole treatments). **H-I**) After an overnight treatment with 250 nM nocodazole the average area of Ribeyeb puncta was significantly lower compared to controls (**H**, P < 0.0001, n = 1008 and 1135 Ribeyeb puncta for control and 250 nM nocodazole treatments). In **I**, the relative frequency of all the areas of Ribeyeb puncta are plotted in nocodazole treatment and controls. For comparisons an unpaired t-test was used in **E-G**, and a Mann-Whitney test was used in **H**. Scale bar in **D** = 5 µm.

We performed a similar analysis on hair cells after a 16-hr treatment with 25 µM taxol. After treatment we observed more acetylated-α-tubulin label, indicating that our taxol treatment successfully stabilized microtubules (qualitative examples: Figure 4-S1A,C, Figure 4-S2A,C). Quantification revealed an overall increase in mean acetylated-α-tubulin label in hair cells after taxol treatment, but this increase did not reach significance (Figure 4-S1E). Unlike nocodazole treatment, taxol did not significantly impact the number of hair cells, the average area of Ribeyeb puncta, or the number of ribbons and precursors (apical and basal) per hair cell compared to controls (Figure 4-S2E,G-I). Interestingly, we observed slightly more complete synapses per hair cell after a 16-hr taxol treatment (Figure 4-S2F). Overall, our long-term taxol treatment indicated that microtubule stabilization did not dramatically impact synapse formation in pLL hair cells.

Together, this pharmacology study revealed that long-term destabilization and stabilization of microtubules during development can impact ribbon formation in pLL hair cells. Microtubule destabilization had the most dramatic effect leading to more ribbons and precursors, and the formation of fewer complete synapses.

### *Kif1aa* mutants have fewer synapses and more ribbon precursors

For cargo (such as ribbons and precursors) to be transported along microtubules, molecular motor proteins are required. Kinesin motor proteins transport cargo along microtubules towards the growing, plus end. Our work demonstrated that in hair cells of the pLL, the plus end of microtubules grow from the apex to the base of the cell (Figure 2). Single-cell RNA sequencing (scRNAseq) has revealed that *kif1aa*, a zebrafish orthologue of mammalian Kif1a, a plus-end directed kinesin motor protein, is highly expressed in pLL hair cells (Lush et al., 2019). In zebrafish, there are 2 orthologues of mammalian *Kif1a*, *kif1aa* and *kif1ab*. ScRNA-seq in zebrafish has demonstrated widespread co-expression of *kif1ab* and *kif1aa* mRNA in the nervous system. Additionally, both scRNA-seq and fluorescent in situ hybridization have revealed that pLL hair cells exclusively express *kif1aa* mRNA (David et al., 2024; Lush et al., 2019; Sur et al., 2023). Therefore, we tested whether Kif1aa could be the kinesin motor that transports ribbons and precursors to the cell base during development.

To test for the role of Kif1aa in pLL hair cells, we created a CRISPR-Cas9 mutant (Varshney et al., 2016). Our *kif1aa* mutant has a stop codon in the motor domain and is predicted to be a null mutation (Figure 5-S1A-B). Recent work in our lab using this mutant has shown that Kif1aa is responsible for enriching glutamate-filled vesicles at the base of hair cells. In addition, this work demonstrated that loss of Kif1aa results in functional defects in mature hair cells including a reduction in evoked post-synaptic calcium responses (David et al., 2024). We hypothesized that Kif1aa may also be playing an earlier role in ribbon formation.

For our initial analysis of *kif1aa* mutants, we co-labeled hair cells at 3 dpf with Ribeyeb to label ribbons and precursors and pan-Maguk to label postsynapses (see examples: Figure 5A-D). This is a similar endpoint used to examine synapse formation after our long-term nocodazole and taxol treatments (Figure 4 and Figure 4-S2). We found that at 3 dpf *kif1aa* mutants had a similar number of hair cells per neuromast (Figure 5E). Despite a similar number of hair cells, we found that there were significantly fewer complete synapses per hair cell in *kif1aa* mutants compared to controls (Figure 5F). In addition, we found that there were significantly more ribbons and precursors (apical and basal) in *kif1aa* mutants compared to controls (Figure 5G-I). As described in the previous section, we also observed fewer complete synapses, and more ribbons and precursors after a 16-hr nocodazole treatment (Figure 4F-G). Together, this suggests that both intact microtubules and Kif1aa are required for normal synapse formation in pLL hair cells.

**Figure 5.**
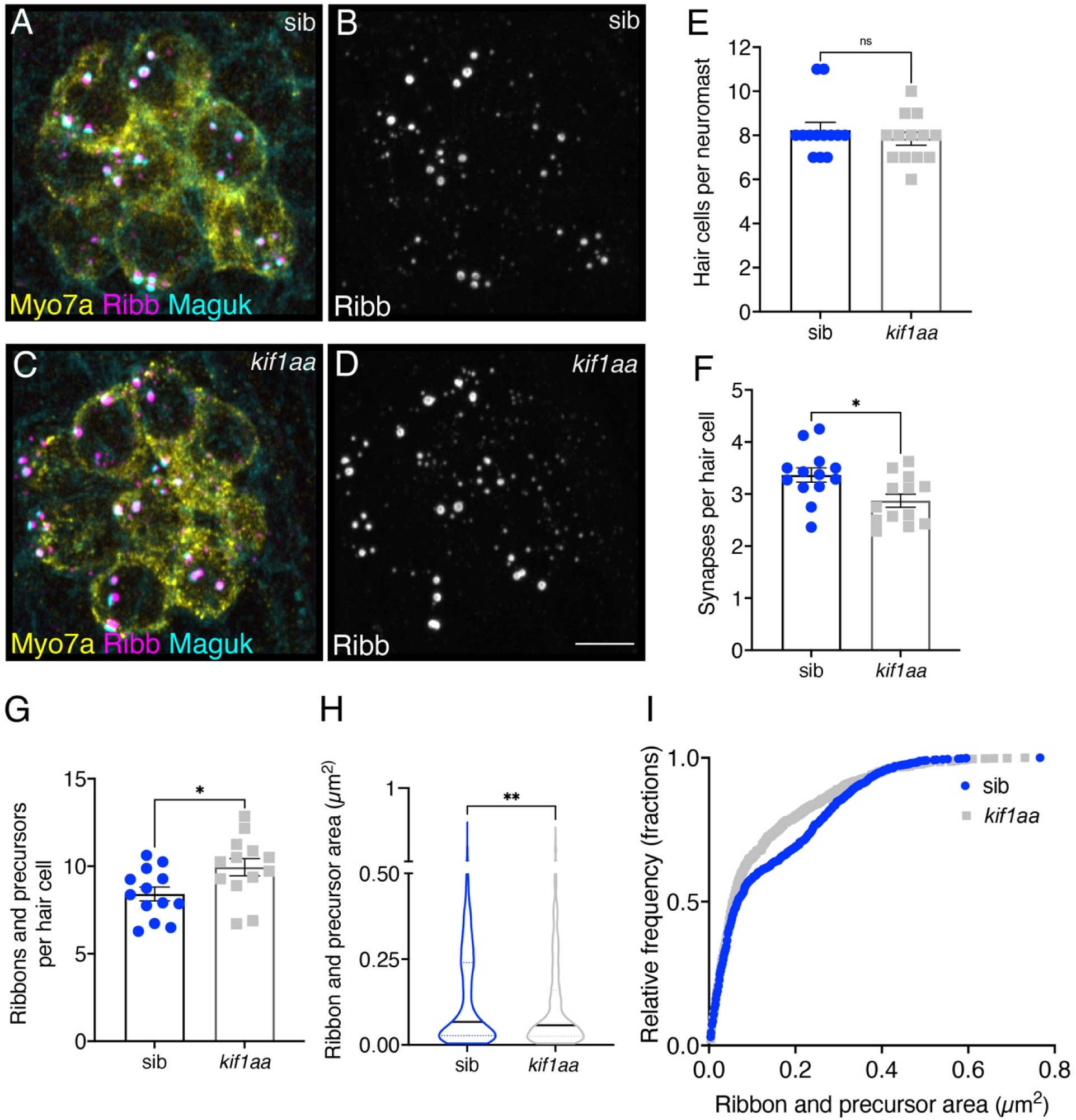
Loss of Kif1aa increases precursor numbers and decreases synapse counts. **A-D)** Example immunostain of neuromasts at 3 dpf in *kif1aa* germline mutants (**C-D**) or sibling control (**A-B**). Myosin7a labels hair cells, Ribeyeb (Ribb) labels precursors and ribbons, and Maguk labels postsynapses. **E-G**) In *kif1aa* mutants there is no change in the number of hair cells per neuromast (**E**, P = 0.418), but there are fewer complete synapses per cell (**F**, P = 0.014), and more ribbons and precursors per cell (**G**, P = 0.024) compared to sibling controls (n = 13 neuromasts for control and *kif1aa* mutants). **H-I**) In *kif1aa* germline mutants the average area of Ribeyeb puncta was significantly lower compared to sibling controls (**H**, P = 0.005, n = 896 and 1008 Ribeyeb puncta for control and *kif1aa* mutants). In **I**, the relative frequency of all the areas of Ribeyeb puncta are plotted in *kif1aa* mutants and sibling controls. For comparisons an unpaired t-test was used in **E-G**, and a Mann-Whitney test was used in **H**. Scale bar in **D** = 5 µm.

### Short-term disruption of microtubules, but not loss of Kif1aa, impacts ribbon formation

Our initial experiments suggest that microtubule networks and Kif1aa are important for proper synapse formation in pLL hair cells. However, the actual changes in precursors and ribbons within developing hair cells was unclear. Therefore, we examined changes in ribbons and precursors in living pLL hair cells over 3-4 hrs of development. For our analysis, we used transgenic lines expressing YFP-tubulin to monitor microtubules and Riba-TagRFP to monitor precursors and ribbons *in vivo*. We examined precursors and ribbons after nocodazole or taxol treatment, or after knockdown of Kif1aa during this developmental window.

We verified the effectiveness of our *in vivo* pharmacological treatments using either 500 nM nocodazole or 25 µM taxol by imaging microtubule dynamics in pLL hair cells (*myo6b:YFP-tubulin*). After a 30-min pharmacological treatment, we used Airyscan confocal microscopy to acquire timelapses of YFP-tubulin (3 µm z-stacks, every 50-100 s for 30-70 min, Movie S8).

Compared to controls, 500 nM nocodazole destabilized microtubules (presence of depolymerized YFP-tubulin in the cytosol, see arrows in Figure 4-S1F-G) and 25 µM taxol dramatically stabilized microtubules (indicated by long, rigid microtubules, see arrowheads in Figure 4-S1F,H) in pLL hair cells. We did still observe a subset of apical microtubules after nocodazole treatment, indicating that this population is particularly stable (see asterisks in Figure 4-S1F-H).

After verifying our *in vivo* pharmacological pharmacology treatments, we acquired Airyscan confocal images of developing hair cells at 2 dpf that express YFP-tubulin and Riba-TagRFP. Then after a 3-4 hr treatment in either 500 nM nocodazole or 25 µM taxol, we reimaged the same hair cells. Consistent with our previous results (Figure 4 and Figure 4-S1,S2) nocodazole and taxol treatments destabilized or stabilized microtubules respectively (qualitative examples: Figure 6A-F). In each developing cell, we quantified the total number of Riba-TagRFP puncta (apical and basal) before and after each treatment. In our control samples, we observed on average no change in the number of Riba-TagRFP puncta per cell (Figure 6G). Interestingly, we observed that nocodazole treatment led to a significant increase in the total number of Riba-TagRFP puncta after 3-4 hrs (Figure 6G). This result is similar to our overnight nocodazole experiments in fixed samples, where we also observed an increase in the number of ribbons and precursors per hair cell. In contrast to our 3-4 hr nocodazole treatment, similar to controls, taxol treatment did not alter the total number of Riba-TagRFP puncta over 3-4 hrs (Figure 6G). Overall, our overnight and 3-4 hr pharmacology experiments demonstrate that microtubule destabilization has a more significant impact on ribbon numbers compared to microtubule stabilization.

**Figure 6.**
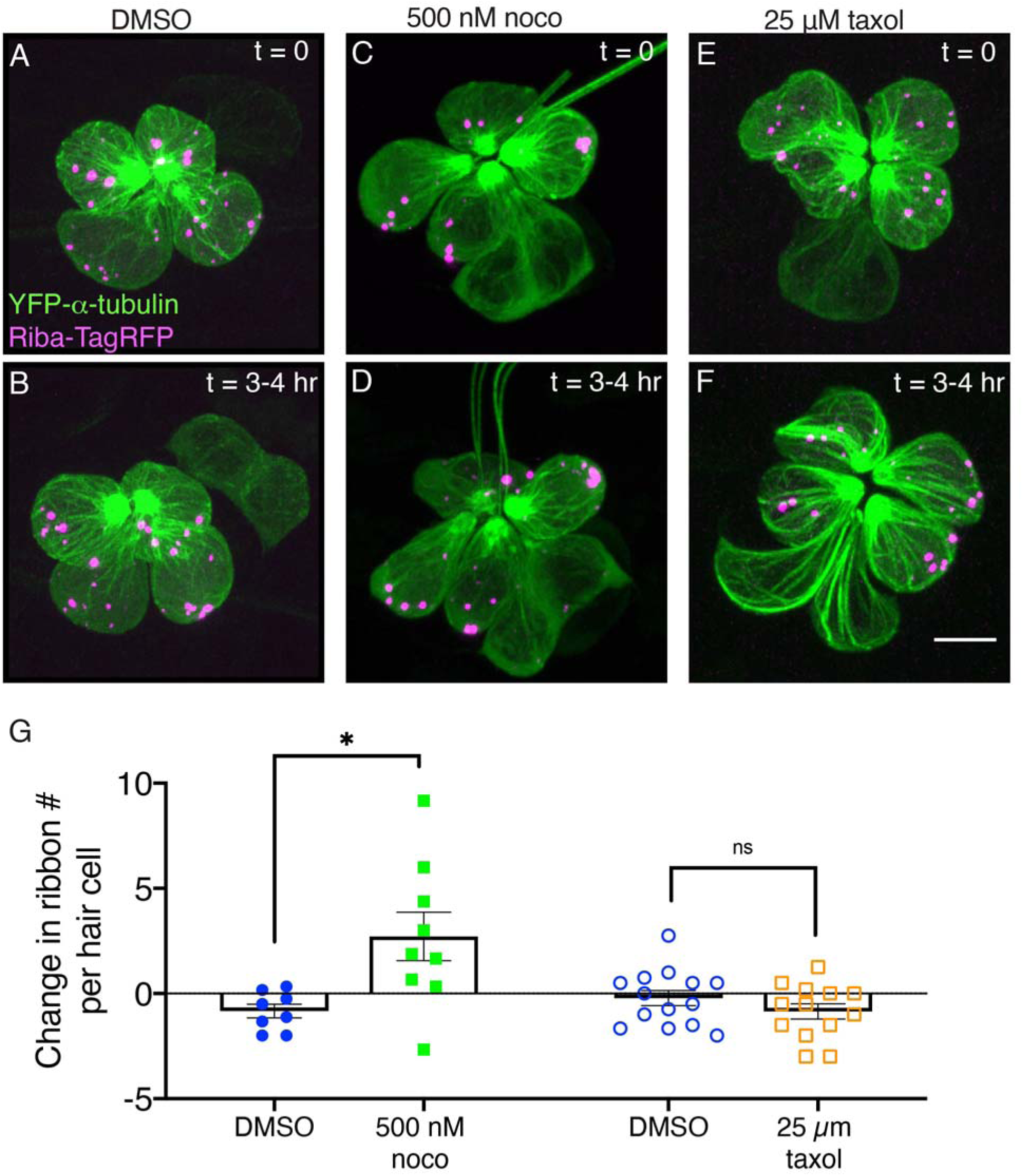
Microtubule destabilization increases ribbon numbers in vivo. **A,C,E)** Example images of neuromasts at 2 dpf. The microtubule network and ribbons are marked with YFP-tubulin and Riba-TagRFP respectively. Neuromasts were imaged immediately after a 30-min treatment with DMSO (**A**, control) 500 nM nocodazole (**C**) or 25 µM taxol (**E**), (t = 0). **B,D,F**) The same neuromasts in **A,C,E** were reimaged after an additional 3-4 hrs of treatment. **G**) Quantification reveals that after 3-4 hr nocodazole treatment, there are more Riba-TagRFP puncta per hair cell compared to controls (n = 9 and 8 neuromasts for nocodazole and DMSO, P = 0.013). In contrast, after a 3-4 hr taxol treatment, there was no significant change in the number of Riba-TagRFP puncta per hair cell compared to controls (n = 13 and 14 neuromasts for taxol and DMSO, P = 0.256). An unpaired t-test was used for comparisons in **G**. Scale bar in **F** = 5 µm.

Next, we used a similar approach to examine the number of Riba-TagRFP puncta in *kif1aa* mutants over a 3-4 hr time window. For this analysis, we examined puncta in *kif1aa* F0 crispants. These mutants are derived from the injection of 2 *kif1aa* guide RNAs (gRNAs) and Cas9 protein. This approach has shown to be an extremely effective way to assay gene function in any genetic background (Hoshijima et al., 2019). Using this approach, we were able to robustly disrupt the *kif1aa* locus (Figure 6-S1). We found that compared to uninjected controls, in *kif1aa* F0 crispants there was no change in the total number of Riba-TagRFP puncta per cell over 3-4 hrs (Figure 6-S2A-F). This result is slightly different than our analysis of *kif1aa* mutants at 3 dpf where we observed significantly more ribbons and precursors per cell compared to controls (Figure 5G). Overall, live imaging over a 3-4 hr time window indicates that a loss of microtubule stability, but not loss of Kif1aa, results in an increase in Riba-TagRFP puncta within developing pLL hair cells.

### Ribbons and precursors require microtubules, but not Kif1aa for directed motion

Our tracking analyses revealed that ribbons and precursors show directed motion on microtubules. Based on these results, we asked what happens to this movement if we disrupt microtubules or the kinesin motor Kif1aa. To test whether microtubules are required for directional ribbon and precursor motion, we used the drugs nocodazole and taxol to alter microtubule dynamics. We treated the fish with these drugs for 30 min and recorded timelapses of ribbon and precursor motion. This short treatment allowed us to observe ribbon motion relatively soon after microtubule disruption and minimize any cytotoxic effects of the compounds. Then we performed MSD and track displacement analyses on ribbon movement to determine if directional motion was impaired (Figure 7A-B).

**Figure 7.**
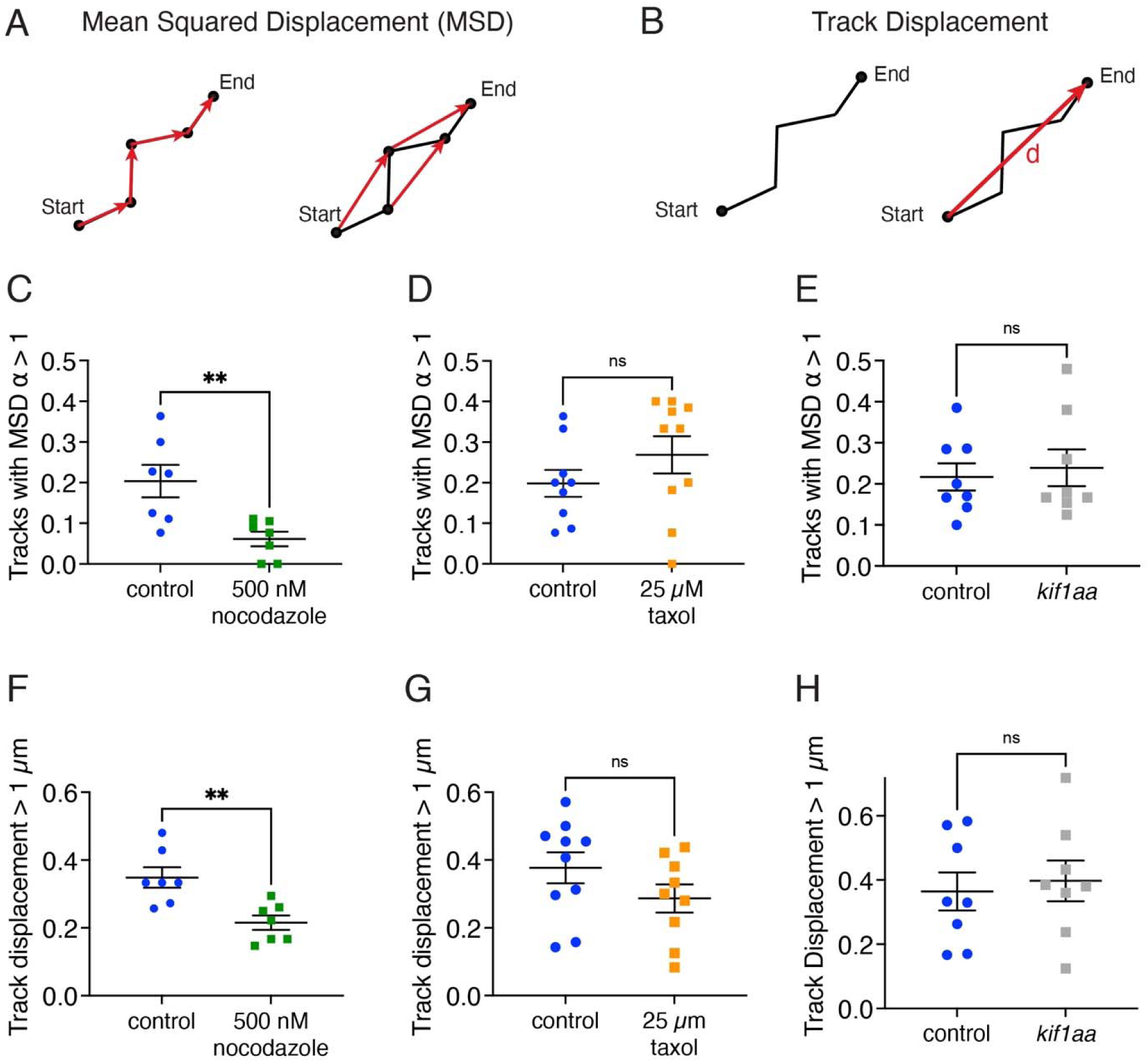
Ribbon precursors require intact microtubules, but not Kif1aa for directional motion. **A** Mean squared displacement (MSD) vs time was used to quantify the proportion of ribbon and precursor tracks with a velocity have α > 1, a behavior indicative of directionally moving tracks. Shown are the first- and second-time steps. **B)** To further quantify directional motion, the proportion of tracks with large displacements >1 µm was quantified. Track displacement was measured as the distance between the start and end point of the track. **C)** Hair cells treated with 500 mM nocodazole have fewer directional tracks (α > 1) compared to controls (P = 0.007). **D)** In hair cells treated with 25 µM taxol there are not significantly fewer directional tracks (α > 1) compared to controls (P = 0.24). **E)** In hair cells lacking Kif1aa, there are not significantly fewer directional tracks (α > 1) compared to controls (P =0.70). **F)** Hairs cells treated with 500 nM nocodazole have fewer ribbons with track displacements > 1 µm compared to control (P = 0.004). **G)** There is no change in track displacement > 1 µm in hair cells treated with 25 µM taxol (P = 0.17). **H)** There is no change in track displacement > 1 µm in hair cells lacking Kif1aa (P = 0.71). N = 7 neuromasts for DMSO control and nocodazole for **C** and **F**; n = 9 and 10 neuromasts for DMSO control and taxol for **D** and **G**; n = 8 neuromasts for control and *kif1aa* for **E** and **H**. An unpaired t-test was used in **C-H**.

Using this approach we found that after destabilizing microtubules by treatment with 500 nM nocodazole for 30 min, the proportion of tracks with α > 1 was significantly reduced compared to controls (Figure 7C). In addition, we also found that the proportion of longer tracks with a cumulative displacement > 1 µm was also reduced after nocodazole treatment compared to controls (Figure 7F). This analysis indicates that an intact microtubule network is needed for proper directional ribbon motion and longer displacements. In contrast, after stabilizing microtubules by treatment with 25 µM taxol, we observed no effect on the proportion of tracks with displacement > 1 µm or the proportion of tracks with α > 1, compared to controls (Figure 7D,G). Interestingly, when we examined the distribution of α values, we observed that taxol treatment shifted the overall distribution towards higher α values (Figure 7-S1A). In addition, when we plotted only tracks with directional motion (α > 1), we found significantly higher α values in hair cells treated with taxol compared to controls (Figure 7-S1B). This indicates that in taxol-treated hair cells, where the microtubule network is stabilized, ribbons with directional motion have higher velocities.

Next, we used a similar approach to examine the tracks of ribbon and precursor motion in *kif1aa* mutants. For this analysis, we tracked ribbons and precursors in *kif1aa* F0 crispants. We observed that the proportion of tracks with displacement > 1 µm was not significantly different between *kif1aa* F0 crispants and controls (Figure 7H). Similarly, the proportion of tracks with α > 1, was also not significantly different from the controls (Figure 7E). Overall, these tracking analyses show that in developing pLL hair cells, the kinesin motor Kif1aa is not essential for the directional motion of ribbons and precursors. In contrast, an unperturbed microtubule network is essential for directed motion.

### Ribbon precursors fuse on microtubules

Our analyses demonstrate that the movement of ribbon precursors along microtubules is important for synapse formation. But in addition to movement, during development it has been proposed that these small precursors come together or fuse to form larger, more mature ribbons (Michanski et al., 2019). This is consistent with our work, where we see a reduction in ribbon and precursor numbers alongside development (Figure 1E). Further, we see an increase in ribbon and precursor number when microtubules are destabilized (Figure 4), indicating an intact microtubule network may also be important for ribbon fusion.

Consistent with this idea, in our timelapses, we observed that ribbons and precursors undergo fusion on microtubules. These fusion events occurred between smaller precursors at the cell apex as well as between larger precursors near the cell base (see examples: Figure 8A-B, Figure 8-S1A-B, Movies S9-S11). We classified an event as a fusion once the ribbons could not be resolved separately and stayed together for the length of the remaining timelapse (or at least 5 min). Although we could not accurately measure the areas of precursors before and after fusion, we did observe that the relative area resulting from the fusion of two smaller precursors was greater than that of either precursor alone. This increase in area suggests that precursor fusion may serve as a mechanism for generating larger ribbons (see examples: Figure 8-S1A-B). Fusion events usually involved ribbon precursors on two separate microtubule filaments coming together during fusion, but fusion events could also occur on the same filament. Although the fusion events were infrequent, within our time windows (30-70 min), we quantified them and counted an average of 1.7 fusions and a maximum of 5 fusions events per neuromast in developing pLL hair cells (n = 27 control neuromasts). Based on the close association with microtubules during these events, we tested whether an intact microtubule network was required for fusion events. We found that after treatment with 500 nM nocodazole for 30 min, there were fewer fusions events (mean 0.4, maximum 2 fusions per neuromast, n = 10 neuromasts). In contrast, the frequency of fusion events remained unchanged upon treatment with 25 µM taxol for 30 min.

**Figure 8.**
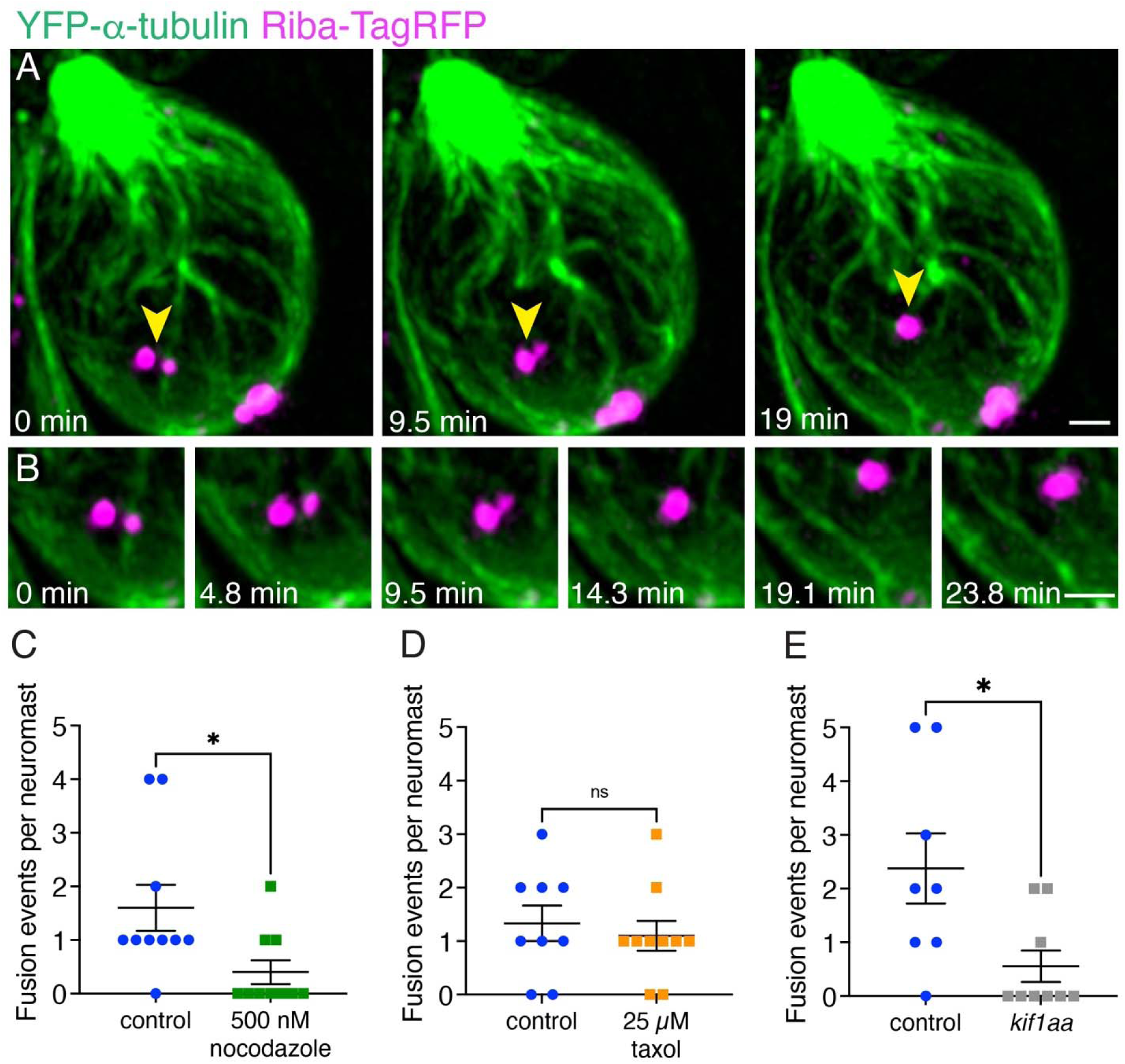
Microtubules and Kif1aa are required for fusion events. **A** An example of two ribbon precursors undergoing fusion on microtubules (yellow arrowheads). **B)** A zoomed-in montage of the example from **A** is shown where the association of each precursor with microtubules can be seen during the process of fusion (also see Movie S9). **C-D)** Destabilizing the microtubules with nocodazole treatment reduces the number of fusion events observed in timelapses (P = 0.023). Taxol treatment has no effect (P = 0.60). **E)** Loss of Kif1aa significantly reduces the number of fusion events observed in timelapses compared to control (P = 0.018). N = 10 neuromasts for DMSO control and nocodazole for **C**; n = 9 and 10 neuromasts for DMSO control and taxol for **D**; n = 8 and 9 neuromasts for control and *kif1aa* for **E**. An unpaired t-test was used for comparisons in **C-E**. Scale bars in **A** and **B** = 1 µm.

In our tracking analysis, we observed that Kif1aa was not required for the directional motion of ribbon and precursors in developing hair cells (Figure 7E,H). However, our immunohistological analyses indicated that there were more precursors and fewer complete synapses in *kif1aa* mutants. Therefore, we examined whether Kif1aa was important for fusion events. For this analysis, we quantified fusion events in *kif1aa* F0 crispants. We observed that similar to nocodazole treatment, there were significantly fewer fusion events in *kif1aa* F0 crispants (mean 0.6, maximum 2 fusions per neuromast, n = 9 neuromasts). This reduction in fusion events may ultimately account for the excess of precursors and fewer complete synapses observed in *kif1aa* mutants (Figure 5). Together our results indicate that during development, ribbon precursors can fuse when associated with microtubules, and an unperturbed microtubule network, along with Kif1aa is needed for these fusion events.

## Discussion

Our work in zebrafish applied high-resolution imaging approaches to investigate how ribbons are assembled and mobilized in developing hair cells. Our study demonstrates that hair cells have microtubule networks that are polarized with growing plus ends pointed to the cell base. Live imaging highlights that during development, ribbons and precursors show directed motion and fusion along microtubules and that an intact microtubule network is important for synapse formation.

### Comparison of ribbon synapse formation in zebrafish and mice

Compared to pLL hair cells (12-18 hrs), synapse formation and refinement occur over a relatively long time window in mouse IHCs (E18-P14). In mice, much of the initial synaptogenesis occurs embryonically, while synapse refinement and pruning occur later, during the first postnatal week. Due to the rapid time course of synapse development in zebrafish, our experiments likely encompass ribbon precursor movement during synaptogenesis and synapse refinement. The companion paper on mouse IHCs focused primarily on ribbon mobility during the dramatic pruning of synapses that occurs during early postnatal development (Voorn et al., 2024). Unfortunately, it was not possible to study earlier events in mouse IHCs, as equivalent experiments were not possible embryonically. Live imaging at postnatal stages in mice indicates that during synapse pruning in IHCs, ribbons may detach from the membrane, undergo local trafficking, and fuse to nearby presynaptic AZs. Together our studies in zebrafish and mouse hair cells illuminate several common features that characterize ribbon and microtubule dynamics during synapse formation.

First, immunostaining in both zebrafish and mice revealed an extensive network of microtubules within hair cells. In both species, the majority of microtubules were stabilized by tubulin acetylation (Figure 2-S1). Our *in vivo* imaging of EB3-GFP in zebrafish hair cells revealed that microtubules are very dynamic, and the plus ends of microtubules grow towards the cell base, towards the presynaptic AZ (Figure 2). This data is consistent with data in mouse IHCs demonstrated via immunostaining that the minus end marker of microtubules, CAMSAP2, localizes to the IHC apex (Vorn et. al.). Overall, both studies demonstrate that microtubule networks in developing hair cells are polarized with the plus ends pointed to the cell base.

Second, live imaging in zebrafish and mouse hair cells revealed that ribbons associate with and move along microtubules. Tracking and quantification of ribbon movement via MSD analysis revealed that in both zebrafish and mouse hair cells, the majority of ribbons show motion indicative of confinement (Figure 3, α < 1). In addition, a subset of ribbon tracks shows evidence of directed motion (Figure 3, α > 1, or displacements > 1 µm). In both species, after using nocodazole to destabilize microtubules, there was a dramatic reduction in instances of directed motion (Figure 7). In addition to ribbon movement, ribbon fusion was observed in the hair cells of mice and zebrafish (Figure 8). Interestingly, in mice many fusion events were balanced out by divisions, or ribbons undergoing separation. In contrast, ribbon divisions were not prevalent in zebrafish hair cells. A lack of ribbon divisions could be due to the rapid speed of development in zebrafish hair cells, where divisions are too transient in nature to be confirmed. Alternatively, divisions may occur primarily late in synapse development and these events were overall less abundant in our zebrafish datasets. Importantly, in both mouse and zebrafish hair cells, ribbon fusion was diminished after microtubule destabilization. Together these results in mice and zebrafish demonstrate that an intact microtubule network is required for the directed motion and fusion of ribbons during development.

Third, we sought to explore the kinesin motor responsible for ribbon movement in hair cells using zebrafish and mouse models. Based on scRNAseq data and previous immunostaining evidence we focused on Kif1a (Lush et al., 2019; Michanski et al., 2019). In both mouse *Kif1a* and zebrafish *kif1aa* mutants, a common observation was an overall reduction in ribbon size (Figure 5). This suggests that a conserved mechanism where both intact microtubules and Kif1aa are required for normal ribbon formation. Despite these results, future work in zebrafish and mice is needed to explore additional kinesin motors that can drive ribbon movement during synapse formation.

### Fusion on microtubules as a mechanism for ribbon enlargement and maturation

Numerous studies have documented small ribbon precursors in developing photoreceptors and hair cells; later in development there are fewer, large ribbons present (Michanski et al., 2019; Regus-Leidig et al., 2009; Schmitz, 2009; Sheets et al., 2011; Sobkowicz et al., 1986, 1982). This has led to the hypothesis that ribbon precursor fusion is a mechanism to form larger, more mature ribbons (Michanski et al., 2019). Our live imaging experiments show that ribbon fusion is indeed a feature of ribbon formation (Figure 8, Figure 8-S1). Interestingly, we observed that fusion often occurs between precursors on adjacent microtubules. Fusion events can occur both apically and basally within hair cells. More apical fusion events are likely a way to increase ribbon size early in development. More basal fusion events could represent the synapse reduction and refinement that occurs later in synapse development. Whether these later, basal fusion events occur after postsynaptic elements are present is unclear and awaits additional studies. Importantly we found that the number of fusion events decreased when microtubule networks were disrupted or when Kif1aa was absent (Figure 8). Together these results confirm the hypothesis that ribbons fuse during development. In addition, our work demonstrates that microtubules and Kif1aa are necessary for fusion and may help facilitate this process.

Ribbons are aggregates of proteins that share many features with biomolecular condensates that form through liquid-liquid phase separation (LLPS) (Wang et al., 2021). Biomolecular condensates are nano-to micro-meter scale compartments that function to concentrate proteins and nucleic acids. Some examples of biomolecular condensates include tight junctions, postsynaptic densities, ribonucleoprotein (RNP) granules, and stress granules (Wang et al., 2021; Wiegand and Hyman, 2020). These condensates are highly mobile and dynamic and constituent molecules diffuse readily and exchange with the surrounding environment. Biomolecular condensates often undergo fusion to form larger ones, and then shrink into a spherical shape (Wiegand and Hyman, 2020). FRAP analyses of Ribeye-GFP labeled ribbons in hair cells have confirmed that tagged Ribeye has a fast recovery phase within ribbons, verifying that Ribeye is highly mobile and dynamic within ribbons (Graydon et al., 2017). Further, these FRAP studies revealed that Ribeye molecules can exchange with the surrounding environment. Our present studies confirmed that ribbon precursors undergo fusion to form larger spherical ribbons (Figure 8A,B). Together these studies point towards the idea that precursors and ribbons are indeed biomolecular condensates. Ribeye also contains sequences predicted to be intrinsically disordered, a feature of molecules that can mediate condensate formation. Interestingly, several biomolecular condensates such as RNP transport granules are known to be transported along the microtubules (Knowles at al., 1996), similar to our observations of ribbon precursor transport.

In our current study, we also found that microtubules are needed for fusion (Figure 8C). In other cellular contexts, the energy released by dynamic microtubules via growth and shrinkage can be used for force generation in a wide range of processes (Vleugel et al., 2016). Interestingly, work on stress granule condensates found a connection between dynamic microtubules and stress granule formation. This work demonstrated that microtubule growth and shrinkage promoted the fusion of small cytoplasmic granules into larger ones (Chernov et al., 2009). In addition, this study also demonstrated that stress granules slide along microtubules and that this movement may act in conjunction with pushing or pulling to promote fusion (Chernov et al., 2009). Ribbons and microtubules may also interact during development to promote fusion. Disrupting microtubules could interfere with this process, preventing ribbon maturation. Consistent with this, short-term (3-4 hr) and long-term (overnight) nocodazole increased ribbon and precursor numbers (Figure 6AG; Figure 4G), suggesting reduced fusion. Long-term treatment (overnight) resulted in a shift toward smaller ribbons (Figure 4H-I), and ultimately fewer complete synapses (Figure 4F).

We also observed that fusion was disrupted in *kif1aa* mutants, despite finding that directed transport of precursors and ribbons remained unchanged in *kif1aa* mutants (Figure 7,8). A recent study on RNP condensates examined kinesin motors and adaptors in the context of microtubules (Cochard et al., 2023). This work found that the precise combination of motor proteins or adaptors can impact where condensates form. Some combinations enabled the formation and transport of condensates along microtubules. However, other combinations restricted condensate formation to microtubule terminals where motor proteins are predicted to form a scaffold. Our work indicates that while more than one kinesin motor protein is capable of transporting ribbon precursors, it is predominantly Kif1aa that mediates fusion along microtubules. This could explain why precursor fusion, but not transport is impacted in *kif1aa* mutants (Figure 7,8). This scenario could also account for the increased number of precursors and the reduced number of complete synapses observed in *kif1aa* mutants (Figure 5).

### Kinesin motors and adaptors for ribbon transport on microtubules

Our immunohistochemistry analyses show clear synaptic defects in *kif1aa* mutants after the bulk of synapse formation has occurred (fewer synapses, more precursors, Figure 5). Despite these defects, we were unable to demonstrate that loss of Kif1aa impacts the movement of precursors along microtubules (Figure 7). These results suggest that another kinesin motor may function to transport precursors and ribbons. Pulldown assays have shown an interaction of Kif3a with Ribeye (Uthaiah and Hudspeth, 2010). While *kif1aa* mRNA is the most abundant kinesin motor protein transcript detected in pLL hair cells, *kif3a* mRNA is also present. *Kif3a* mRNA is present at lower levels and is expressed more broadly in pLL hair cells, supporting cells, and afferent neurons (Sur et al., 2023). Therefore, both Kif1aa and Kif3a may be competent to transport developing ribbons. Future work exploring the role of Kif3a separately or in combination with Kif1aa will illuminate the role these motors play in synapse assembly in hair cells. In addition, it will be useful to visualize these kinesins by fluorescently tagging them in live hair cells to observe whether they associate with ribbons.

Regardless of the anterograde kinesin motor(s) that transports precursors to the cell base, we also observed that precursors can also move in the retrograde direction, towards the cell apex (Figure 3C bottom panels, Figure 3L, and Figure 3-S1B). In fact, out of the tracks with directional motion, we found that 20.8 % of precursors moved apically. This could be due to the fact that a subpopulation of microtubules shows plus-end mediated growth apically (22.8 %).

Alternatively, it is possible that retrograde motors may also transport ribbon precursors. For example, this could be accomplished using minus-end directed motors, such as cytoplasmic dynein heavy chain, or kinesins in the kinesin-14 family (Sweeney and Holzbaur, 2018; Yamada et al., 2017). In the future, it will be important to explore the role of these additional motor proteins in the context of ribbon and precursor mobility.

Our findings indicate that ribbons and precursors show directed motion indicative of motor-mediated transport (Figure 3 and 7). While a subset of ribbons moves directionally with α values > 1, canonical motor-driven transport in other systems, such as axonal transport, can achieve even higher α values approaching 2 (Bellotti et al., 2021; Corradi et al., 2020). We suggest that relatively lower α values arise from the highly dynamic nature of microtubules in hair cells. In axons, microtubules form stable, linear tracks that allow kinesins to transport cargo with high velocity. In contrast, the microtubule network in hair cells is highly dynamic, particularly near the cell base. Within a single time frame (50-100 s), we observe continuous movement and branching of these networks. This dynamic behavior adds complexity to ribbon motion, leading to frequent stalling, filament switching, and reversals in direction. As a result, ribbon transport appears less directional than the movement of traditional motor cargoes along stable axonal filaments, resulting in lower α values compared to canonical motor-mediated transport. Notably, treatment with taxol, which stabilizes microtubules, increased α values to levels closer to those observed in canonical motor-driven transport (Figure 7-S1). This finding supports the idea that the relatively lower α values in hair cells are a consequence of a more dynamic microtubule network. Overall, this dynamic network gives rise to a slower, non-canonical mode of transport.

Another important component of motor-mediated transport is adaptor proteins that selectively link cargo to specific motor proteins (Fu and Holzbaur, 2014). In neurons, cargos leave the Golgi apparatus and are packed into specialized transport vesicles, along with adaptor scaffolds (Farías et al., 2012; Sampo et al., 2003). Work in neurons has shown that the Kif1a motor transports Rab3-positive synaptic vesicle precursors (Okada et al., 1995). This transport is thought to rely on DENN (differentially expressed in normal and neoplastic cells)/MADD (MAP kinase activating death domain) which links synaptic vesicle precursors to Kif1a (Hummel and Hoogenraad, 2021; Niwa et al., 2008). Interestingly, like mature ribbons, ribbon precursors are decorated with synaptic vesicles (Michanski et al., 2019). Thus, is it possible that these synaptic vesicles may contain adaptor proteins that couple the ribbon to a kinesin motor to enable transport. In the future, it will be important to identify this adaptor protein and to understand what other molecules are co-transported with Ribeye during ribbon formation.

### Ribbon formation in the absence of microtubules

Our work indicates that over short time scales (30-70 min), microtubule destabilization via nocodazole treatment reduces the number of ribbons and precursors that show directed motion indicative of active transport (Figure 7). In addition, prolonged nocodazole treatment throughout development (16 hrs) leads to the formation of fewer synapses (Figure 4). But in both treatment paradigms, some ribbons still show directed motion, and some synapses continue to form despite microtubule disruption. If tracks of microtubules are required for developing precursor and ribbon mobility during development, what underlies this residual movement and synapse formation?

One possibility is that not all microtubules are disrupted after nocodazole treatment. While high doses of nocodazole (∼40 µM) eliminate all microtubules, they are also cytotoxic (Laisne et al., 2021). The doses used in our experiments (100-500 nM) were not cytotoxic, although higher doses did result in the death of hair cells. At these lower doses, microtubules— especially the more apical, stable population (see examples: Figure 2-S1B, Figure 4-S1F-G)— were not entirely disrupted. Residual ribbon mobility during nocodazole treatment could occur along these remaining microtubules. Alternatively, other cytoskeletal components, such as actin or intermediate filaments may contribute to ribbon mobility. Work in mouse IHCs has shown that actin filaments help regulate and organize synaptic vesicles at mature ribbon synapses (Guillet et al., 2016). In addition, in mice, the actin-based motor protein Myosin6 has been implicated in ribbon-synapse formation and function (Roux et al., 2009). A third possibility is that ribbons and precursors move through the cytoplasm via diffusion and are captured at the AZ via an adapter protein. Although we primarily observed ribbons and precursors attached to microtubules, when these filaments are destabilized using nocodazole, movement could still occur via diffusion. Once near the base of the cell, presynaptic molecules such as Bassoon, could act to anchor ribbons at the AZ (Jing et al., 2013). Any of these scenarios—residual microtubule-based transport, alternative cytoskeletal component, or diffusion followed by capture—could explain how ribbons continue to move and how synapses still form in the absence of intact microtubules. In all likelihood, a combination of these processes is required to ensure that ribbon synapses form properly. Additional pharmacological and imaging experiments are needed to delineate the relative contributions of these processes.

Another important consideration is the potential off-target effects of nocodazole. Even at non-cytotoxic doses, nocodazole toxicity may impact ribbons and synapses independently of its effects on microtubules. While this is less of a concern in the short- and medium-term experiments (30 min to 4 hr), long-term treatments (16 hrs) could introduce confounding effects. Additionally, nocodazole treatment is not hair cell-specific and could disrupt microtubule organization within afferent terminals as well. Thus, the reduction in ribbon-synapse formation following prolonged nocodazole treatment may result from microtubule disruption in hair cells, afferent terminals, or a combination of the two.

### Does spontaneous activity shape ribbon transport or fusion?

In previous work, we demonstrated that spontaneous calcium activity impacts ribbon formation in pLL hair cells (Sheets et al., 2012; Wong et al., 2019). Spontaneous activity is well documented in developing sensory systems [reviewed in: (Leighton and Lohmann, 2016)]. In the inner ear of mammals, spontaneous activity in sensory hair cells is thought to act during development to establish neuronal connections within the inner ear, and downstream in the brain to shape tonotopic maps (Ceriani et al., 2019; Tritsch et al., 2007). In our previous work in pLL hair cells, we found that spontaneous rises in presynaptic calcium loads calcium into synaptic mitochondria. Calcium loading into the mitochondria regulates the amount of NAD^+^ to NAD(H) in the developing hair cell. Blocking either presynaptic or mitochondria calcium during development results in higher levels of NAD^+^, the formation of larger ribbons, fewer synapses, and the retention of small ribbon precursors in hair cells (Sheets et al., 2012; Wong et al., 2019). What aspect of ribbon formation is impacted by spontaneous activity remains unclear.

Work in yeast has shown that calcium is important for microtubule stability (Adamíková et al., 2004; Façanha et al., 2002). In addition, work in dendritic spines has shown that synaptic calcium responses promote microtubule entry into active spines (Merriam et al., 2013).

Therefore, it is possible that spontaneous calcium activity may act to stabilize microtubules or facilitate movement to the presynaptic AZ to facilitate ribbon transport. In addition to altering the microtubule network, spontaneous activity could also impact ribbon fusion. Protein condensate formation and fusion are influenced by many aspects of the cellular environment, including temperature, pH, osmolarity, and ion concentration (Wang et al., 2022). Elevated calcium can promote the fusion of chromogranin proteins that undergo LLPS in the Golgi lumen (Gerdes et al., 1989; Yoo, 1995). Therefore, it is possible that elevated calcium during a spontaneous calcium event could promote ribbon or precursor fusion. In addition to the cellular environment, posttranslational modifications to the proteins within the condensate can impact formation and LLPS. Such modifications include: phosphorylation, acetylation, SUMOylation, ubiquitination, methylation, and ADP-ribosylation (Luo et al., 2021). These modifications can alter protein-protein interactions by changing the charge, structure, or hydrophobicity. Our previous work indicated that NAD^+^ or NAD(H) via a NAD^+^/NADH binding domain on Ribeye impacts ribbon formation (Wong et al., 2019). It is possible that in addition to calcium, NAD^+^/NADH levels in the cell could modify Ribeye to facilitate condensate formation or fusion. In the future, it will be important to use the live imaging approaches outlined here to understand how spontaneous presynaptic- and mitochondrial-calcium influx impact the movement and fusion of ribbon precursors.

Overall, our live imaging studies demonstrate that microtubule networks are critical for ribbon and precursor mobility, and fusion in developing zebrafish hair cells. However, ribbon synapses contain many molecules, and synapse formation requires many successive steps. In the future it will be important to develop approaches to endogenously tag and label other presynaptic (Ca_V_1.3 channels, Bassoon, Piccolino) and postsynaptic proteins (PSD95, GluR2/3/4) that make up ribbon synapses. For this future work, zebrafish is an ideal model system for creating these new genetic tools and for live imaging. By imaging ribbons, along with these other tagged synaptic components, we will gain a more comprehensive picture of synapse formation. Understanding how ribbon synapses form is essential to determining how to reform synapses when they are disrupted in auditory and visual disorders.

## Methods

### Zebrafish animals

Zebrafish (*Danio rerio*) were bred and cared for at the National Institutes of Health (NIH) under animal study protocol #1362-13. Zebrafish larvae were raised at 28°C in E3 embryo medium (5ILmM NaCl, 0.17 mM KCl, 0.33ILmM CaCl_2_, and 0.33 mM MgSO_4_, buffered in HEPES, pH 7.2). All experiments were performed on larvae aged 2-5 days post fertilization (dpf). Larvae were chosen at random at an age where sex determination is not possible. The previously described mutant and transgenic lines were used in this study: *Tg(myo6b:ctbp2a-TagRFP)*^idc11Tg^ referred to as *myo6b:riba-TagRFP; Tg(myo6b:YFP-Hsa.TUBA)*^idc16Tg^ referred to as *myo6b:YFP-tubulin* (Ohta et al., 2020; Wong et al., 2019). *Tg(myo6b:ctbp2a-TagRFP)*^idc11Tg^ reliably labels mature ribbons, similar to a pan-CTBP immunolabel at 5 dpf (Figure 1-S1A-B). This transgenic line does not alter the number of hair cells or complete synapses per hair cell (Figure 1-S1A-D). In addition, *myo6b:ctbp2a-TagRFP* does not alter the size of ribbons (Figure 1-S1E).

### Zebrafish transgenic and CRISPR-Cas9 mutant generation

To create *myo6b:EB3-GFP* transgenic fish, plasmid construction was based on the tol2/gateway zebrafish kit (Kwan et al., 2007). The p5E *pmyo6b* entry clone (Trapani et al., 2009) was used to drive expression in hair cells. A pME-*EB3-GFP* clone was kindly provided by Catherine Drerup at the University of Wisconsin, Madison. pDestTol2pACryGFP was a gift from Joachim Berger & Peter Currie (Addgene plasmid # 64022). These clones were used along with the following tol2 kit gateway clone, p3E-*polyA* (#302) to create the expression construct: *myo6b:EB3-GFP*. To generate the stable transgenic fish line *myo6b:EB3-GFP*^idc23Tg^, plasmid DNA and tol2 transposase mRNA were injected into zebrafish embryos as previously described (Kwan et al., 2007). The *myo6b:EB3-GFP*^idc23Tg^ transgenic line was selected for a single copy and low expression of EB3-GFP.

A *kif1aa* germline mutant (*kif1aa*^idc24^) was generated using CRISPR-Cas9 technology as previously described (Varshney et al., 2016). Exon 6, containing part of the Kinesin motor domain was targeted (Figure 5-S1A). Guides RNAs (gRNAs) targeted to *kif1aa* are as follows: 5’-ACGGATGTTCTCGCACACGT(AGG)-3’, 5’-GTGCGAGAACATCCGTTGCT(AGG)-3’, 5’-TGGACTCCGGGAATAAGGCT(AGG)-3’, 5’-AGAATACCTAGCCTTATTCC(CGG)-3’. Founder fish were identified using fragment analysis of fluorescent PCR (fPCR) products. A founder fish containing a complex insertion or deletion (INDEL) that destroys a BslI restriction site in exon 6 was selected (Figure 5-S1B). This INDEL disrupts the protein at amino acid 166 (Figure 5-S1A).

Subsequent genotyping was accomplished using standard PCR with touchdown, and BslI restriction enzyme digestion. *Kif1aa* genotyping primers used were: *kif1aa*_FWD 5’-AACACCAAGCTGACCAGTGC-3’ and *kif1aa*_REV 5’-TGCGGTCCTAGGCTTACAAT-3’.

Because *kif1aa* mutants have no phenotype to distinguish them from sibling controls at the ages imaged, the low throughput of our live imaging approaches made using germline mutants prohibitive. Therefore, we created *kif1aa* F0 crispants for our live imaging analyses.

Here we injected the following *kif1aa* gRNAs: 5’-GTGCGAGAACATCCGTTGCT(AGG)-3’ and 5’-AGAATACCTAGCCTTATTCC(CGG)-3’, along with Cas9 protein, as previously described (Hoshijima et al., 2019). We then grew *kif1aa* injected F0 crispants for 2 days and then used them for our live imaging analyses. Studies have shown that F0 crispants are a fast and effective way to knock down gene function in any genetic background (Hoshijima et al., 2019; Sheets et al., 2021). After live imaging, we genotyped all *kif1aa* F0 crispants (Figure 6-S1) to ensure that the gRNAs cut the target robustly using fPCR and the following primers: *kif1aa*_FWD_fPCR 5’-TGTAAAACGACGGCCAGT-AAATAGAGATTCACTTTTAATC-3’ and *kif1aa*_REV_fPCR 5’-GTGTCTT-CCTAGGCTTACAATGCTTTTGG-3’ (Carrington et al., 2015). fPCR fragments were run on a genetic analyzer (Applied Biosystems, 3500XL) using LIZ500 (Applied Biosystems, 4322682) as a dye standard. Analysis of fPCR revealed an average peak height of 4740 a.u. in wild type, and an average peak height of 126 a.u. in *kif1aa* F0 crispants (Figure 6-S1E-F). Any *kif1aa* F0 crispant without robust genomic cutting or a peak height > 500 a.u. was not included in our analyses.

### Zebrafish pharmacology

To destabilize or stabilize microtubules, larval zebrafish at 2 dpf were incubated in either nocodazole (Sigma-Aldrich, SML1665) or Paclitaxel (taxol) (Sigma-Aldrich, 5082270001). Both drugs were maintained in DMSO. For experiments these drugs were diluted in media for a final concentration of 0.1 % DMSO, 250-500 nM nocodazole, and 25 µM taxol. For controls, larvae were incubated in media containing 0.1 % DMSO. For long-term incubation (16 hr), wild-type larvae were incubated in E3 media containing 250 nM nocodazole or 25 µM taxol at 54 hpf for 16 hrs (overnight). After this long-term treatment, larvae were fixed and prepared for immunohistochemistry (see below). For live, short-term incubations (for 3-4 hr incubations or ribbon tracking), transgenic larvae (*myo6b:riba-tagRFP; Tg(myo6b:YFP-alpha-tubulin*) at 48-54 hpf were embedded in 1 % low melt agarose prepared in E3 media containing 0.03 % tricaine (Sigma-Aldrich, A5040, ethyl 3-aminobenzoate methanesulfonate salt). 500 nM nocodazole, 25 µM taxol, or DMSO were added to the agarose and to the E3 media used to hydrate the sample. For short-term treatments, hair cells were imaged after 30 min of embedding.

### Immunohistochemistry of zebrafish samples

Immunohistochemistry used to label acetylated-α-tubulin, tyrosinated-α-tubulin, Ribeyeb or pan-CTBP (ribbons and precursors), pan-Maguk (postsynaptic densities), and Myosin7a (cell bodies) was performed on whole zebrafish larvae similar to previous work. The following primary antibodies were used: rabbit anti-Myosin7a (Proteus 25-6790; 1:1000); mouse anti-pan-Maguk (IgG1) (Millipore MABN7; 1:500); mouse anti-Ribeyeb (IgG2a) ((Sheets et al., 2011); 1:10,000); mouse anti-CTPB (IgG2a) (Santa Cruz sc-55502; 1:1000); mouse anti-acetylated-α-tubulin (IgG2b) (Sigma-Aldrich T7451; 1:5,000); mouse anti-tyrosinated-α-tubulin (IgG2a) (Sigma-Aldrich MAB1864-I; 1:1,000); chicken anti-GFP (to stain YFP-tubulin) (Aves labs GFP-1010; 1:1,000). The following secondary antibodies were used at 1:1,000: (ThermoFisher Scientific, A-11008; A-21143, A-21131, A-21240; A-11-039, A-21242, A-21241, A-11039). Larvae were fixed with 4 % paraformaldehyde in PBS for 4ILhr at 4°C. All wash, block, and antibody solutions were prepared in 0.1 % Tween in PBS (PBST). After fixation, larvae were washed 5 × 5 min in PBST. Prior to block, larvae were permeabilized with acetone. For this permeabilization larvae were first washed for 5 min with H_2_O. The H_2_O was removed and replaced with ice-cold acetone and samples were placed at −20°C for 3 min, followed by a 5 min H_2_O wash. The larvae were then washed for 5 × 5 min in PBST. Larvae were then blocked overnight at 4°C in blocking solution (2 % goat serum, 1 % bovine serum albumin, and 2 % fish skin gelatin in PBST). Larvae were then incubated in primary antibodies in antibody solution (1 % bovine serum albumin in PBST) overnight, nutating at 4°C. The next day, the larvae were washed for 5 × 5 min in PBST to remove the primary antibodies. Secondary antibodies in antibody solution were added and larvae were incubated for 3 hrs at room temperature. After 5IL×IL5ILmin washes min in PBST to remove the secondary antibodies, larvae were rinsed in H_2_O and mounted in Prolong gold (ThermoFisher Scientific, P36930).

### Confocal imaging and analysis of fixed zebrafish samples

After immunostaining, fixed zebrafish samples were imaged on an inverted Zeiss LSM 780 (Zen 2.3 SP1) or an upright Zeiss LSM 980 (Zen 3.4) laser-scanning confocal microscope with Airyscan using a 63x 1.4 NA oil objective lens. Z-stacks encompassing the entire neuromast were acquired every 0.17 (LSM 980) or 0.18 (LSM 780) µm with an 0.04 µm x-y pixel size and Airyscan autoprocessed in 3D.

Synaptic images from fixed samples were further processed using FIJI. Acetylated-α-tubulin or Myosin7 label was used to manually count hair cells. Complete synapses comprised of both a Ribeyeb/CTBP and Maguk puncta were also counted manually. To quantify the area of each ribbon and precursor, images were processed in FIJI using a macro, ‘IJMacro_AIRYSCAN_simple3dSeg_ribbons only.ijm’ as previously described (Wong et al., 2019). Here each Airyscan z-stack was max-projected, and background corrected using rolling-ball subtraction. A threshold was applied to each image, followed by segmentation to delineate individual Ribeyeb/CTBP puncta. The watershed function was used to separate adjacent puncta. A list of 2D objects of individual ROIs (minimum size filter of 0.002 μm^2^) was created to measure the 2D areas of each Ribeyeb/CTBP puncta. Areas for all Ribeyeb/CTBP puncta within each neuromast were then exported as a csv spreadsheet. Areas were averaged per neuromast, per genotype or plotted in a frequency distribution. For comparisons, all fixed images analyzed in FIJI were imaged and processed using the same parameters.

To quantify the mean intensity of acetylated-α-tubulin after overnight nocodazole or taxol treatments, 20 slices centered on the hair cells were max-projected in FIJI. An ROI was drawn around the hair cells, and this ROI was used to measure the mean intensity of the acetylated-α-tubulin label in each neuromast.

### Confocal imaging and in vivo analysis of ribbon numbers in developing zebrafish hair cells

For counting ribbon numbers in developing and mature hair cells (Figure 1), double transgenic *myo6b:riba-TagRFP* and *myo6b:YFP-tubulin* larvae at 2 and 3 dpf were imaged. Transgenic larvae were pinned to a Sylgard-filled petri dish in E3 media containing 0.03 % tricaine and imaged on a Nikon A1R upright confocal microscope using a 60x 1 NA water objective lens. Denoised images were acquired using NIS Elements AR 5.20.02 with an 0.425 µm z-interval, at, 16x averaging, and 0.05 µm/pixel. Z-stacks of whole neuromasts including the kinocilium were acquired in a top-down configuration using 488 and 561 nm lasers. The 488 nm laser along with a transmitted PMT (T-PMT) detector was used to capture the kinocilial heights.

For the quantification of ribbon numbers at different developmental stages (Figure 1), a custom-written Fiji macro “Live ribbon counter” was used to batch-process the z-stacks. The red channel (Riba-TagRFP) of each z-stack was thresholded (threshold value = 97). Watershed was applied to the thresholded stack to separate ribbons near each other. The resulting mask from the thresholding and watershedding was applied to the original red channel. The number of ribbons was then counted using ‘3D Objects Counter’ plugin (Threshold = 1, min size = 0, max size = 183500). The counted objects were merged with the green channel (YFP-tubulin). Each z-stack was visually inspected to determine the localization of the ribbons. Ribbons below the nucleus were classified as ‘basal’ and the rest as ‘apical’. The number of apical and basal ribbons was counted in each hair cell.

To classify the developmental stage of each hair cell (Figure 1), the height of the kinocilium was used. The number of z-slices between the kinocilium tip and base was determined and multiplied by the z-slice interval (0.425 µm) to get the kinocilium height. Hair cells with heights < 1.5 µm were classified as ‘early’, hair cells with heights 1.5-10 µm were classified as ‘intermediate’ and hair cells with heights 10-18 µm were classified as ‘late’. Hair cells with heights > 18 µm were considered ‘mature’.

### Confocal imaging and **in vivo** tracking EB3-GFP dynamics in zebrafish

Transgenic *myo6b:EB3-GFP* larvae at 2-3 dpf were mounted in 1 % LMP agarose containing 0.03 % tricaine in a glass-bottom dish. Larvae were imaged on an inverted Zeiss LSM 780 (Zen 2.3 SP1) confocal microscope using a 63x 1.4 NA oil objective lens. For timelapses, confocal z-stacks of partial cell volumes (3.5 µm, 7 z slices at 0.5 µm z interval) with an 0.07 µm x-y pixel size were taken every 7 s for 15-30 min.

The EB3-GFP timelapses were registered in FIJI using the plugin “Correct 3D drift” (Parslow et al., 2014), max-projected, and then tracked in 2D in Imaris. For spot detection, we used an estimated xy diameter of 0.534 µm with background subtraction. The detected spots were filtered by ‘Quality’ using the automatic threshold. The timelapses were visually checked to make sure the spot detection was accurate. For the tracking step, the ‘Autoregressive motion’ algorithm was used, with a maximum linking distance of 1 µm and a maximum gap size of 3 frames. To ensure accurate track detection, short tracks were removed by filtering for the number of spots in a track (> 5) and track displacement length (> automatic threshold).

To calculate the track angles relative to the cell base, we used cells that lie horizontally so we only need to consider the angles in the 2D, xy plane. In Imaris, the tracks in each hair cell were selected and exported separately. Using the start and end position coordinates of the exported tracks, we calculated track angles in MATLAB using custom-written code called, “EB3 track angle”. The angle of each hair cell was measured in Imaris. The final track angle distribution plotted was obtained by measuring the difference between each track angle and the angle of the hair cell.

To create movies of EB3-GFP tracks in Movie S1, the FIJI plugin “Correct 3D drift” was applied to the timelapse. Z-stacks were then max-projected, and tracks were detected using the FIJI plugin TrackMate (Parslow et al., 2014; Tinevez et al., 2017). For Movie S1, the LoG detector in TrackMate was used with an estimated object diameter of 0.6 µm, and a quality threshold of 8, using a median filter and sub-pixel localization. The Linear Assignment Problem (LAP) tracker was selected using a frame to frame linking max distance of 1 µm, a track segment gap closing max distance of 1 µm and a max frame gap of 2 µm. Tracks were colored by Track index. For viewing tracks over time, tracks were displayed as “Show tracks backwards in time” with a fade range of 5-time points. To create a color-coded temporal map of EB3-GFP tracks over a short time window (21 s, Figure 2C-D), the FIJI Hyperstack plugin “Temporal-Color code” was used with the 16 colors LUT.

### Confocal imaging and **in vivo** analysis of ribbon numbers after short-term pharmacological treatments in zebrafish

For counting ribbons after 3-4 hr drug treatment, transgenic zebrafish expressing *myo6b:riba-TagRFP* and *myo6b:YFP-tubulin* at 2 dpf were examined. Transgenic larvae were mounted in 1 % low melt agarose in E3 media containing 0.03 % tricaine and one of the following: 500 nM nocodazole, 25 µM taxol or 0.1 % DMSO (control). Samples were imaged on an inverted Zeiss LSM 780 (Zen 2.3 SP1) confocal microscope with Airyscan, along with a 63x NA 1.4 oil objective lens. Z-stacks encompassing the entire neuromast were acquired every 0.18 µm with an 0.04 µm x-y pixel size and Airyscan autoprocessed in 3D.

To quantification of ribbon numbers before and after 3-4 hr nocodazole and taxol treatment or in *kif1aa* F0 crispants (Figure 6, Figure 6-S2), the custom-written Fiji macro “Live ribbon counter” described above was used to batch-processed the z-stacks. The red channel (Riba-TagRFP) of each z-stack was thresholded (threshold value = 28) and segmented (watershed). The resulting mask was applied to the original red channel. The number of ribbons was then counted using ‘3D Objects Counter’ (threshold = 1, min size = 0, max size = 183500).

The counted objects were merged with the green channel (YFP-tubulin). Each z-stack was visually inspected to make sure the objects counted were within hair cells. The number of ribbons per neuromast was determined and divided by the number of hair cells. The difference in ribbon numbers pre- and post-drug treatment was plotted.

### Confocal imaging and ***in vivo*** tracking of ribbons

To visualize ribbon precursor movement, timelapses of double transgenic *myo6b:riba-TagRFP* and *myo6b:YFP-tubulin* larvae at 2 dpf were imaged. For pharmacological treatments, transgenic larvae were mounted in 1 % low melt agarose in E3 media containing 0.03 % tricaine and one of the following: 500 nM nocodazole, 25 µM taxol or 0.1 % DMSO (control) in a glass-bottom dish. Double transgenic *kif1aa* crispants and uninjected controls were mounted in 1 % low melt agarose in E3 media containing 0.03 % tricaine in a glass-bottom dish. Larvae were imaged on an inverted Zeiss LSM 780 or an upright LSM 980 confocal microscope with Airyscan using a 63x 1.4 NA oil objective lens. Airyscan z-stacks of partial cell volumes (∼3 µm, 15-20 z-slices using 0.18 µm z-interval and an 0.04 µm x-y pixel size) were taken on the LSM 780 every 50-100 s for 30-70 min. Faster LSM 980 Airyscan z-stacks of partial cell volumes (∼2-3.5 µm, 12-20 z-slices using 0.17 µm z interval and an 0.04 µm x-y pixel size) were taken every 3-20 s for 5-40 min. Airyscan timelapses were autoprocessed in 3D. In addition, we acquired a subset of LSM 780 Airyscan z-stacks every 5-8 min for 30-100 min to capture fusion events more clearly for Movies S9-11.

Timelapses were registered using the FIJI plugin “Correct 3D drift”. Drift-corrected timelapses were then tracked in 3D in Imaris using spot detection with estimated xy diameters of 0.427 µm (with background subtraction). The spots were filtered based on ‘Quality’, with thresholds between 3-8, chosen after visual inspection of the detected spots. For tracking, the ‘Autoregressive motion’ algorithm was used with a maximum linking distance of 1.13 µm and a maximum gap size of 1 frame. Tracks with number of spots < 5 were not included. Using the track displacement length filter in Imaris, the number of tracks with track displacement length > 1 µm were counted and divided by the total number of tracks to get the fractions plotted in Figure 3 and 7. Vectors generated in Imaris were used to manually determine the location and direction of tracks with track displacement length > 1 µm. For the mean squared displacement (MSD) analysis, the xyzt coordinates were exported in ‘csv’ format for all tracks in a timelapse. The MSD analysis was done using the prewritten MATLAB class MSDanalyzer (Tarantino et al., 2014). MSDanalyzer calculates the mean squared displacement for each track, curve-fits the MSD vs time, and provides the value of the exponent (a). The first 25 % of the MSD vs time graph was used for curve-fitting. Tracks with the number of spots < 10 were removed to ensure the accuracy of the MSD analysis. Fusion events between ribbons and precursors were scored manually in these timelapses.

### Statistics

All data shown are mean ± standard error of the mean (SEM). All experiments were compiled from data acquired on at least two independent days from different clutches. All replicates were biological–distinct animals and cells. Wild-type animals were selected at random for drug treatments. Datasets were excluded if there was excessive x-y-or z drift. In all datasets dot plots represent the ‘n’. N represents either the number of neuromasts, hair cells, synapses, or puncta as stated in the legends. For all zebrafish experiments, a minimum of 3 animals and 6 neuromasts were examined. Sample sizes were selected to avoid Type 2 error. All statistical analyses were performed using Prism 10 software (GraphPad). A D’Agostino-Pearson normality test was used to test for normal distributions. To test for statistical significance between two samples, either unpaired t-tests (normally distributed data) or Wilcoxon or Mann-Whitney tests (non-normally distributed data) were used. For multiple comparisons, a one-way ANOVA was used. A P value less than 0.05 was considered significant.

## Contributions

KK, SH, and H-TW did the immunohistochemistry, along with the imaging and analysis of fixed samples. KP and MU created the *kif1aa* mutants. KP created *kif1aa* crispants for analyses and made the *myo6b:EB3-GFP* transgenic line. SH did all the live imaging, along with the quantifications and tracking analyses. KK and SH made figures and wrote the manuscript.

## Data Availability Statement

All data and code used in this paper are available on Dryad: DOI: 10.5061/dryad.crjdfn3gg.

## Acknowledgements

We thank Drs Juan Angueyra and Katie Drerup for their comments on our manuscript. This work was supported by the National Institute on Deafness and Other Communication Disorders (NIDCD) Intramural Research Program Grant 1ZIADC000085-01 (KK) and Project B08 of the Collaborative Research Center 889 ‘*Cellular Mechanisms of Sensory Processing*’ of the German Research Foundation (MU, awarded to Christian Vogl).

## Supplemental Figures

**Figure 1-S1.**
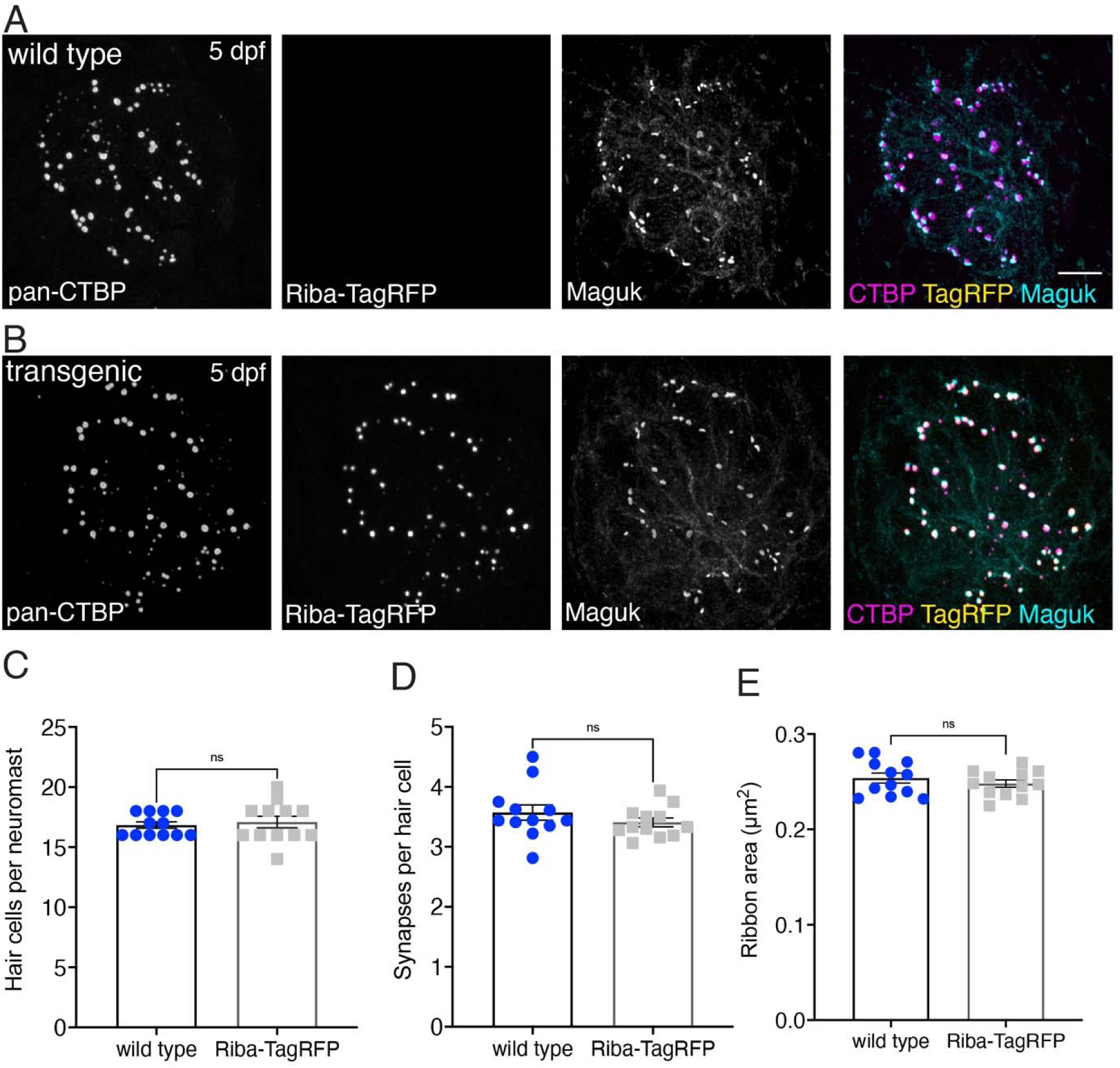
Riba-TagRFP transgenic fish have normal cell numbers, synapses per hair cell, and ribbon areas. **A-B)** Example immunostained images from pLL hair cells of wild type **(A)** and Riba-TagRFP transgenic fish **(B)** are shown at 5 dpf. The neuromasts are labeled with pan-CTBP (labels ribbons) and Maguk (labels postsynapses). **C-E)** Quantification of these images shows no significant differences in the number of hair cells (**C**, P = 0.728), the number of synapses per hair cell (**D**, P = 0.283), or the ribbon areas > 0.1 µm^2^ (**E**, P = 0.392) between the wild type and transgenic (n = 12 neuromasts for wild type and Riba-TagRFP). For comparisons a Mann-Whitey test was used in C and an unpaired t-test was used in **D-E**, scale bar in **A** = 5 µm.

**Figure 1-S2.**
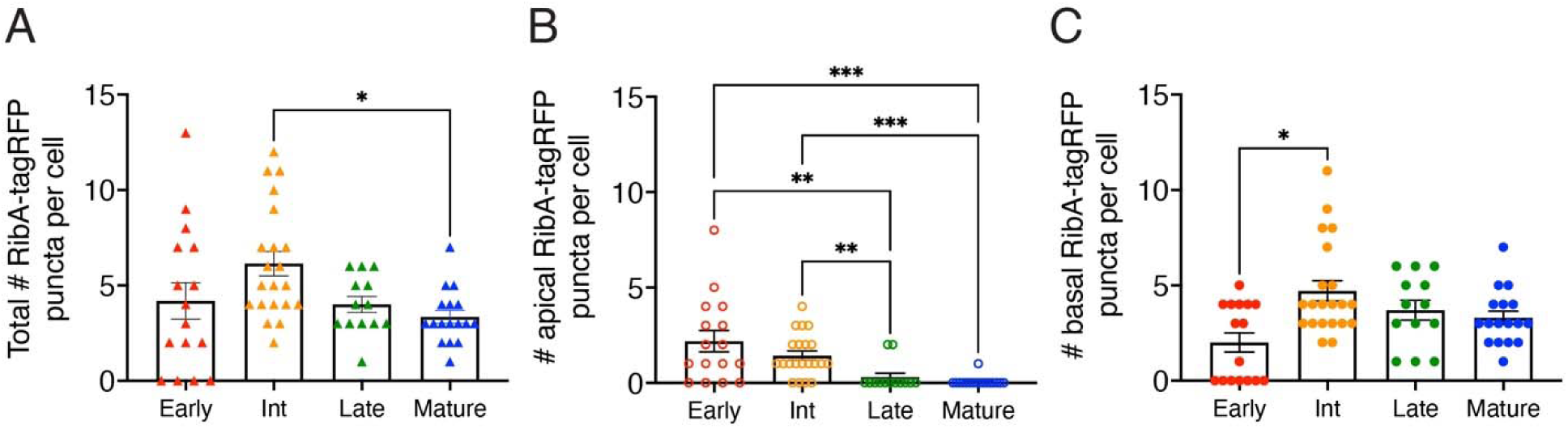
Ribbon number and apical-basal localization changes during development. **A** The total number of Riba-TagRFP puncta increases from early to intermediate stages. The total number of puncta becomes significantly reduced upon maturation. **B)** The number of apically-localized Riba-TagRFP precursors is high at early and intermediate stages and is significantly reduced compared to late and mature hair cells. **C)** The number of basally-localized Riba-TagRFP ribbons is low at early stages and becomes significantly higher by intermediate stages. For comparisons, a Kruskal-Wallis test was used in **A-C**, * P < 0.05, ** P < 0.01, *** P < 0.001.

**Figure 2-S1.**
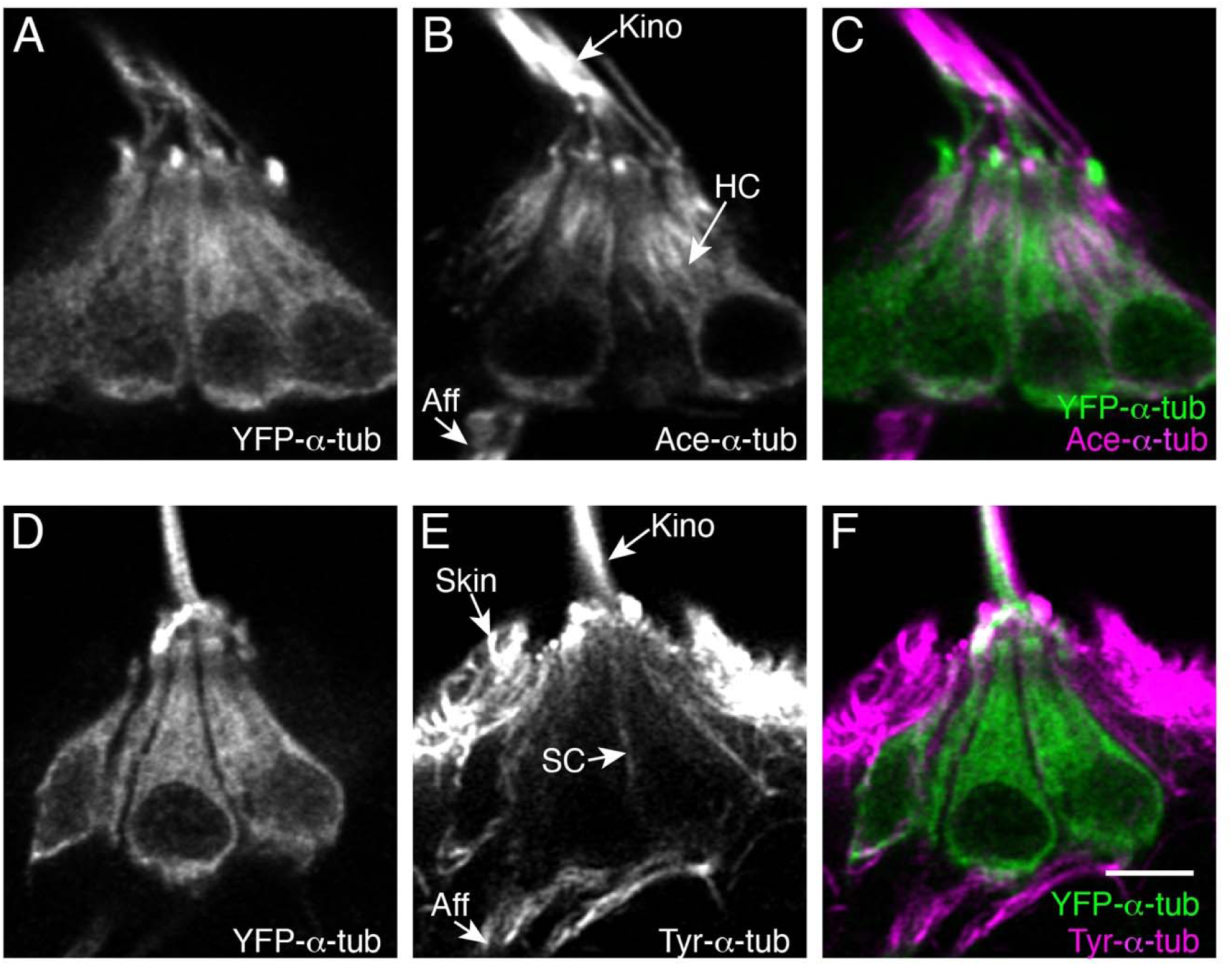
Microtubule modifications in lateral-line hair cells. **A-F)** Example immunostains of pLL hair cells expressing YFP-α-tubulin at 5 dpf, labeled either with acetylated-α-tubulin (**A-C**) or tyrosinated-α-tubulin (**D-F**). Panels **A** and **D** show YFP-α-tubulin, **B** and **E** show acetylated-α-tubulin or tyrosinated-α-tubulin labels respectively. Panels **C** and **F** show the merged images. HC = hair cell; SC = supporting cell; Aff = afferent process; Kino = kinocilium; Scale bar in **F** = 5 µm.

**Figure 2-S2.**
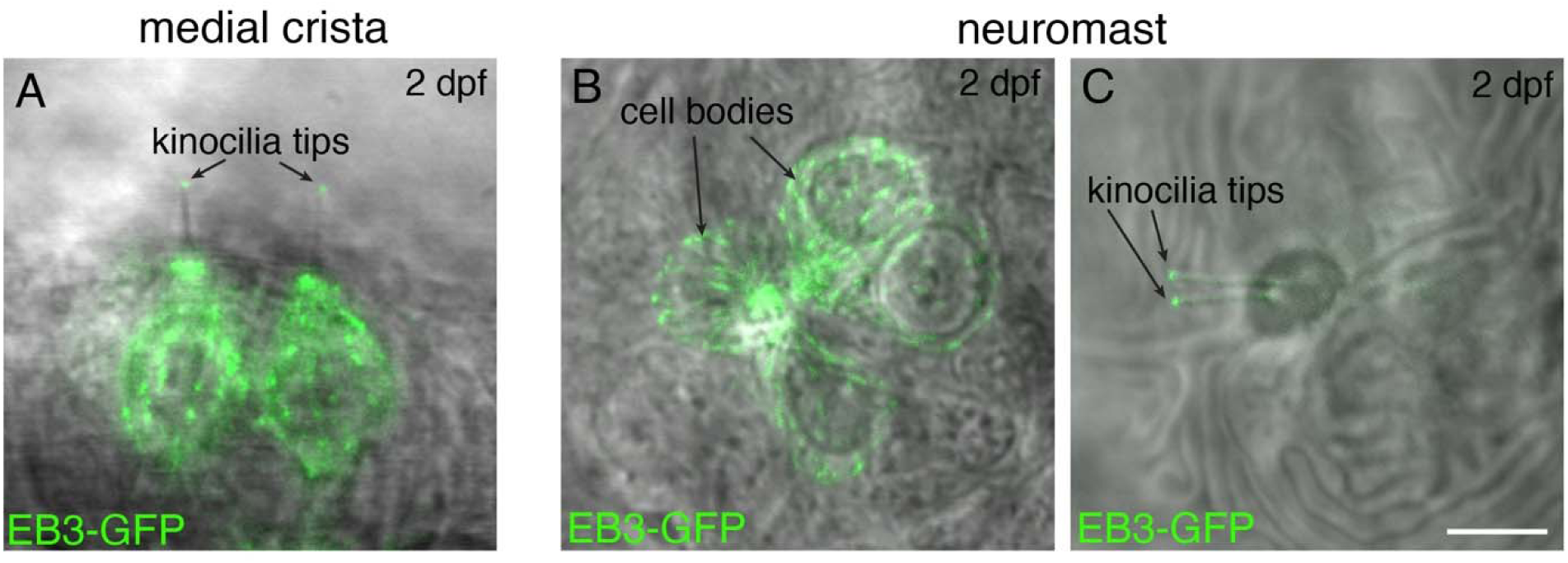
EB3-GFP label reveals microtubule (+) ends at kinocilial tips in hair cells. **A** Example side-view image of EB3-GFP localization in 2 hair cells of the medial cristae within the zebrafish inner ear at 2 dpf. **B-C)** Example images of EB3-GFP localization in 4 hair cells within a pLL neuromast at 2 dpf. In **B**, a maximum-intensity projection of EB3-GFP localization in the cell bodies is shown. In **C**, a more apical projection shows the localization of EB3-GFP in the kinocilia of the same hair cells as **B**. In both zebrafish inner ear and pLL hair cells EB3-GFP is present in the cell bodies and at the tips of kinocilia. In all images EB3-GFP is overlaid onto a transmitted light image. Scale bar in **C** = 5 µm.

**Figure 3-S1.**
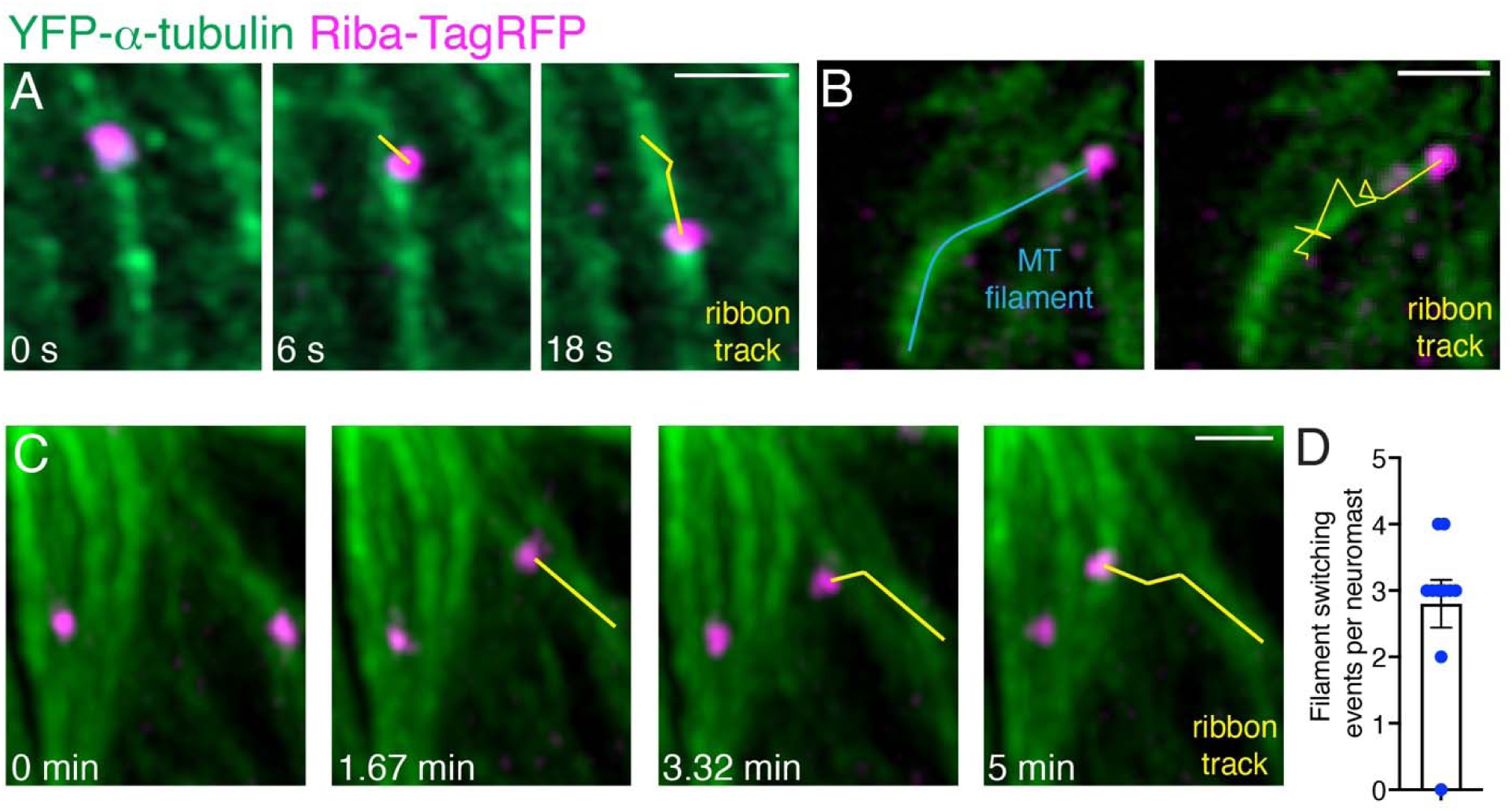
Ribbons move directionally on microtubules and can move between microtubule filaments. **A** Example of ribbon movement on a microtubule directed towards the cell base. Ribbon precursor movement was imaged at 3 s intervals for ∼5 min. Shown in **A** are frames 10, 12 and 16 (also see Movie S5). **B)** Example of a ribbon movement along a microtubule towards the cell apex. Images were acquired every 50 s for ∼40 min. The microtubule (MT) filament is indicated in blue. The track of the ribbon over frames 25-46 (21 frames, 17.5 min) is shown in yellow. The track is overlaid onto the last image of the timelapse (also see Movie S6). **C)** Example of a ribbon moving along a microtubule towards the cell apex and then switching to another microtubule. Images were acquired every 98 s for 43 min. Shown in **C** are frames 10, 12, 14, and 16 (also see Movie S7). **D)** Quantification shows an average of 2.8 filament switching events per neuromast during timelapses (timelapses acquired every 50-100 s for 30-70 min, n = 10 neuromasts). Scale bars in **A-C** = 1 µm.

**Figure 4-S1.**
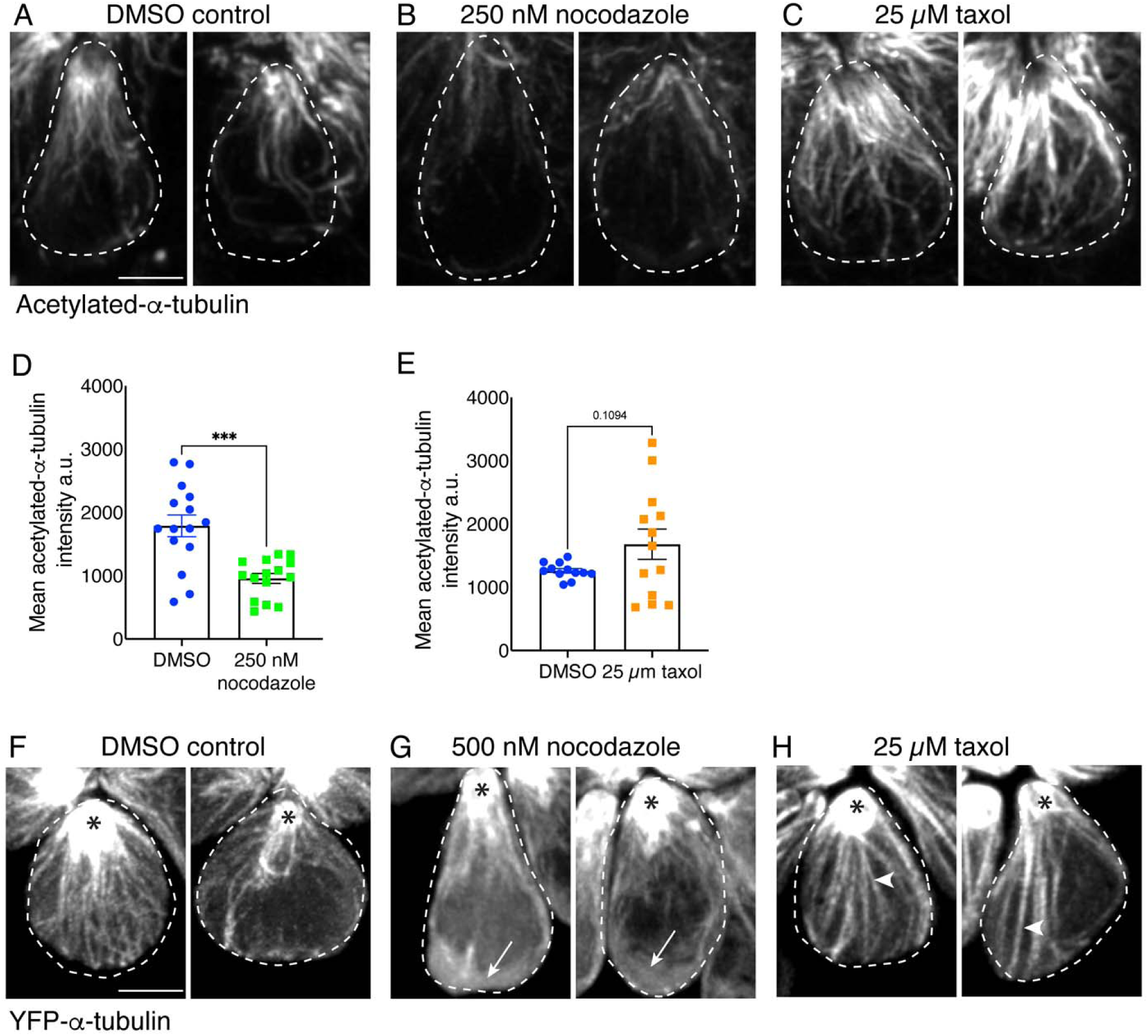
Nocodazole and taxol treatment impact microtubules in lateral line hair cells. **A-C)** Example side-view images of individual hair cells treated overnight with 250 nM nocodazole **(B)** or 25 µM taxol **(C)** overnight compared to DMSO control **(A)**. Hair cells were fixed and stained with acetylated-α-tubulin at 3 dpf. In **B**, fewer microtubule networks are observed in nocodazole-treated hair cells. In contrast, in **C**, more intense, stable microtubules are observed in taxol-treated hair cells. **D)** Quantification reveals a significant reduction in the mean acetylated-α-tubulin intensity in hair cells treated overnight with nocodazole, indicating microtubule disruption (n = 15 neuromasts for each condition, P < 0001). **E)** Although the mean acetylated-α-tubulin intensity levels were elevated after treatment with 25 µM taxol overnight, the elevation was not significant (n = 12 neuromasts for DMSO control and 13 for taxol treatment, P = 0.109). **F-H)** Example side-view images of individual hair cells at 2 dpf expressing YFP-α-tubulin treated at 2 dpf with 500 nM nocodazole **(G)** or 25 µM taxol **(H)** for 3-4 hrs, compared to DMSO controls **(F)**. In **G** fewer microtubule networks along with more diffuse YFP-α-tubulin label (white arrows) is observed in nocodazole-treated hair cells. In contrast, in **H**, more intense and long, stable microtubules (white arrowheads) are observed in taxol-treated hair cells. All hair cells in **F-H** show persistent and stable microtubules at the cell apex (black asterisks). Images in **A-C** and **F-H** were maximum-intensity projected and displayed using the same settings. For comparisons, an unpaired t-test was used in **D** and **E.** Scale bar in **A** and **F** = 2.5 µm.

**Figure 4-S2.**
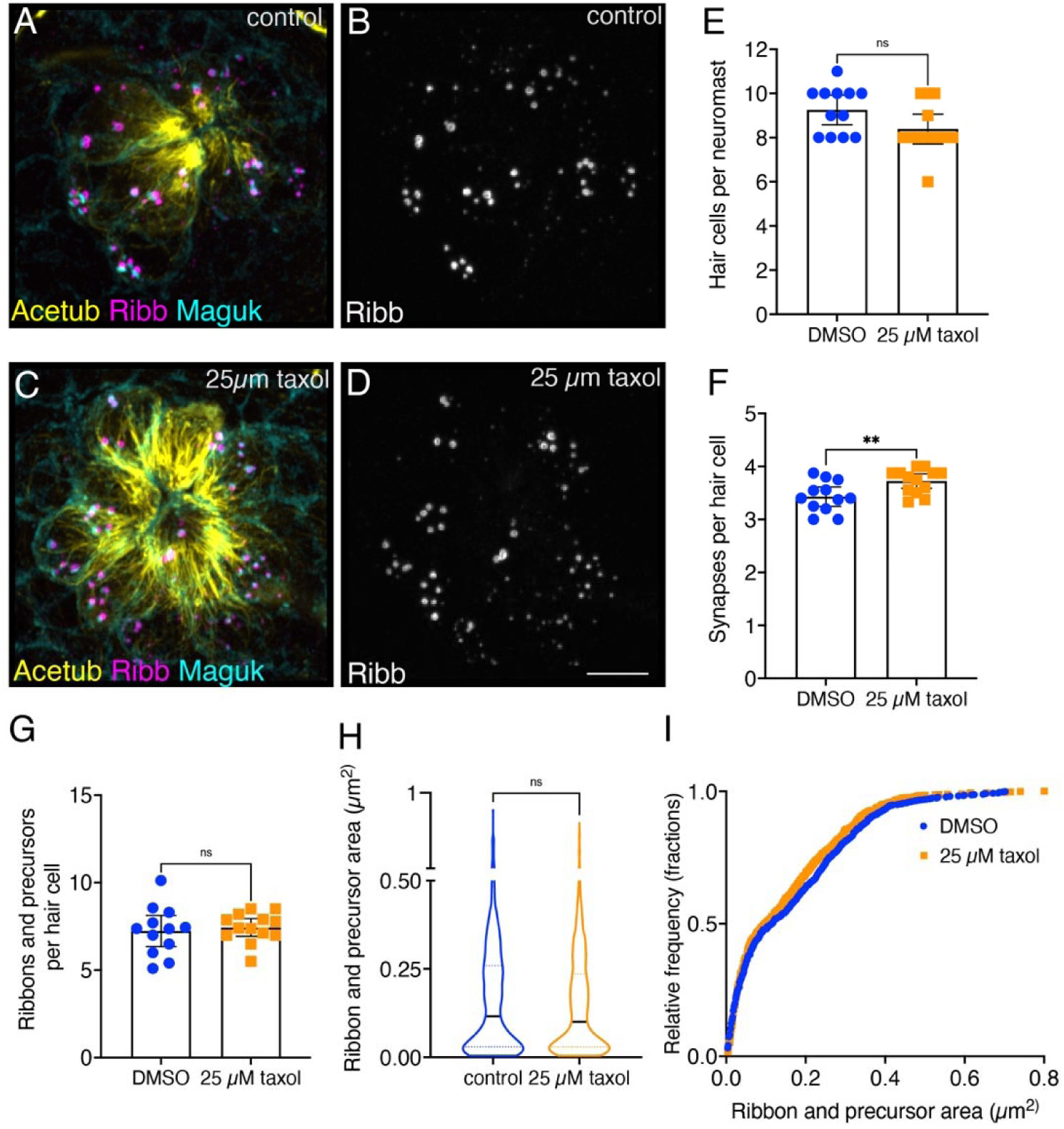
Overnight microtubule stabilization slightly increases synapse counts. **A-D)** Example immunostain of a neuromast at 3 dpf after an overnight treatment with 25 µM taxol (**C-D**) or DMSO (**A-B**). Acetylated-α-tubulin (Acetub) labels microtubules, Ribeyeb (Ribb) labels precursors and ribbons, and Maguk labels postsynapses. **E-G**) After an overnight treatment with 25 µM taxol there are similar numbers of hair cells per neuromast (**E**, P = 0.059), more complete synapses per cell (**F**, P = 0.009), and no change in the number of ribbons and precursors per cell (**G**, P = 0.672) compared to controls (n = 12 and 13 neuromasts for control and 25 µM taxol treatments). **H-I**) After an overnight treatment with 25 µM taxol the average area of Ribb puncta was not changed compared to controls (**H**, P = 0.153, n = 800 and 817 Ribb puncta for control and 25 µM taxol treatments). In **I**, the relative frequency of all the areas of Ribb puncta are plotted in taxol treatment and controls. For comparisons an unpaired t-test was used in **E-G**, and a Mann-Whitney test was used in **H**. Scale bar in **D** = 5 µm.

**Figure 5-S1.**
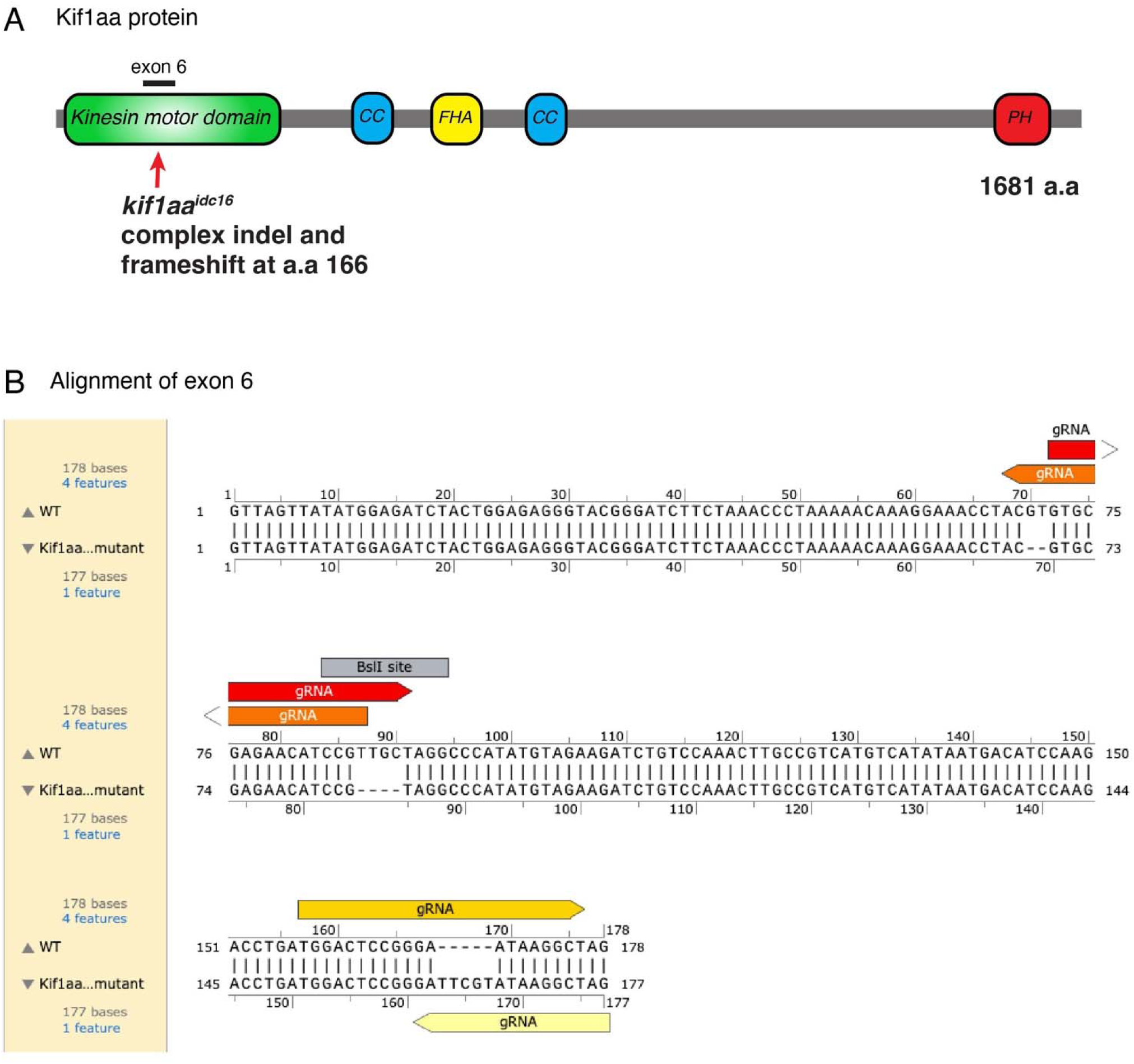
Kif1aa protein and exon 6 lesions. **A** Overview of the Kif1aa protein and major domains (coiled coil (CC), fork-head associated (FHA), pleckstrin homology (PH)). The location of the germline *kif1aa* lesion in the kinesin motor domain within exon 6 is indicated. **B)** The DNA sequence of exon 6 (178 bp) in wild type and *kif1aa* germline mutants. The 4 gRNAs used to make the *kif1aa* germline mutant are shown. 2 deletions and 1 insertion are present in *kif1aa* germline mutants. The BslI restriction site used for genotyping is shown. This DNA alignment was done in Snapgene.

**Figure 6-S1.**
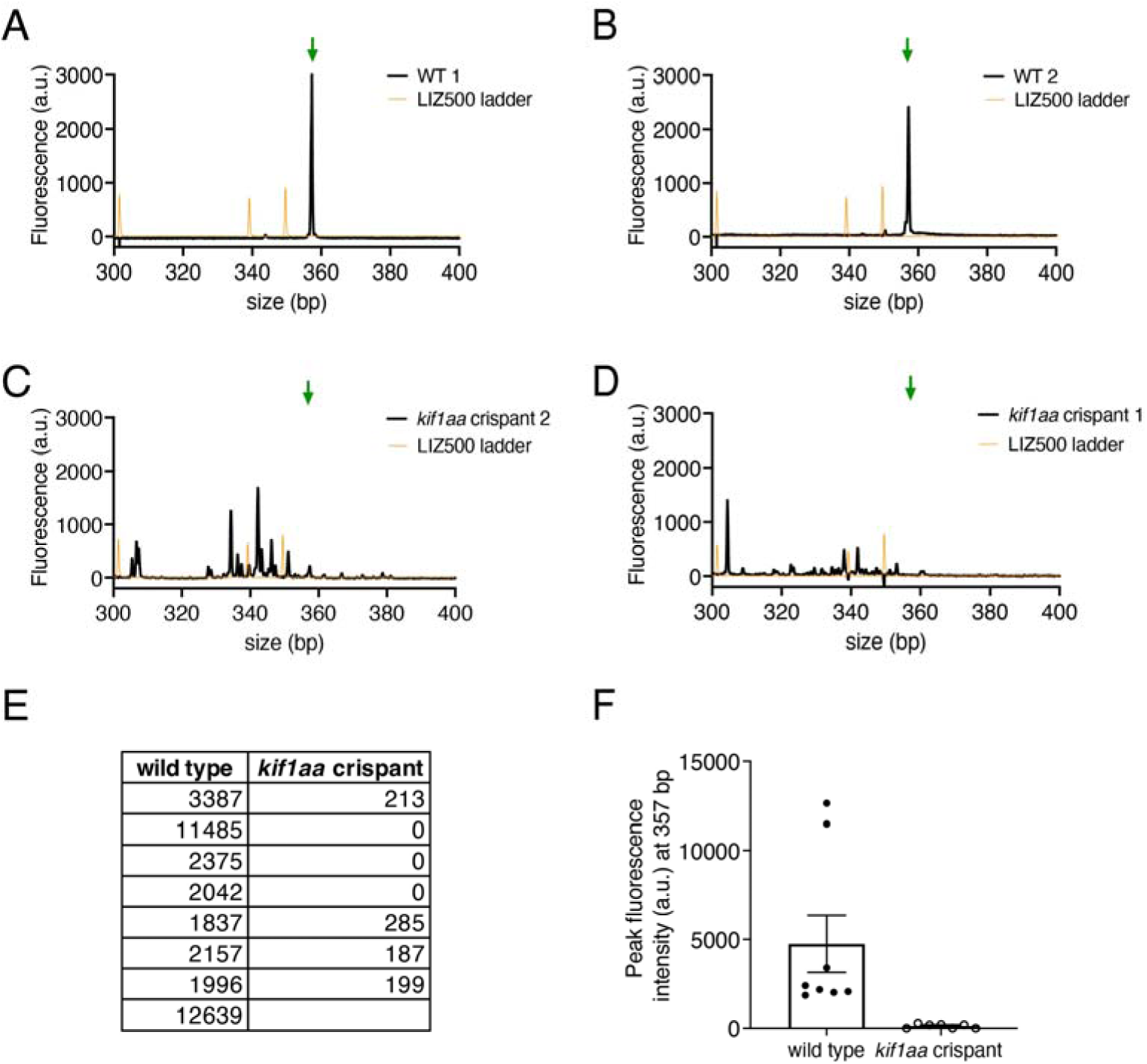
*kif1aa* crispant verification via genotyping via fluorescent fragment analysis. Each fish that was imaged in our kif1aa experiments was genotyped using fragment analysis of fluorescent PCR products (primers: kif1aa_FWD_fPCR 5’-TGTAAAACGACGGCCAGT-AAATAGAGATTCACTTTTAATC-3’ and kif1aa_REV_fPCR 5’-GTGTCTT-CCTAGGCTTACAATGCTTTTGG-3’ (Carrington et al., 2015)). **A-D**) Shown are example graphs from wild type (**A,B**) and kif1aa crispants (**C,D**). The wild-type graphs have a distinct peak at 357 bp (green arrow) while the kif1aa crispants this peak is dramatically reduced. The multiple secondary peaks in **C-D** are indicative of indels up to ∼50 bp in kif1aa crispants. **E-F**) Quantification of fluorescence intensity values (a.u.) at 357 bp for wild type and kif1aa crispant are shown in a table (**E**) and a graph (**F**), demonstrating successful genomic cutting in the kif1aa crispants. kif1aa crispants without robust genomic cutting were excluded from our analyses.

**Figure 6-S2.**
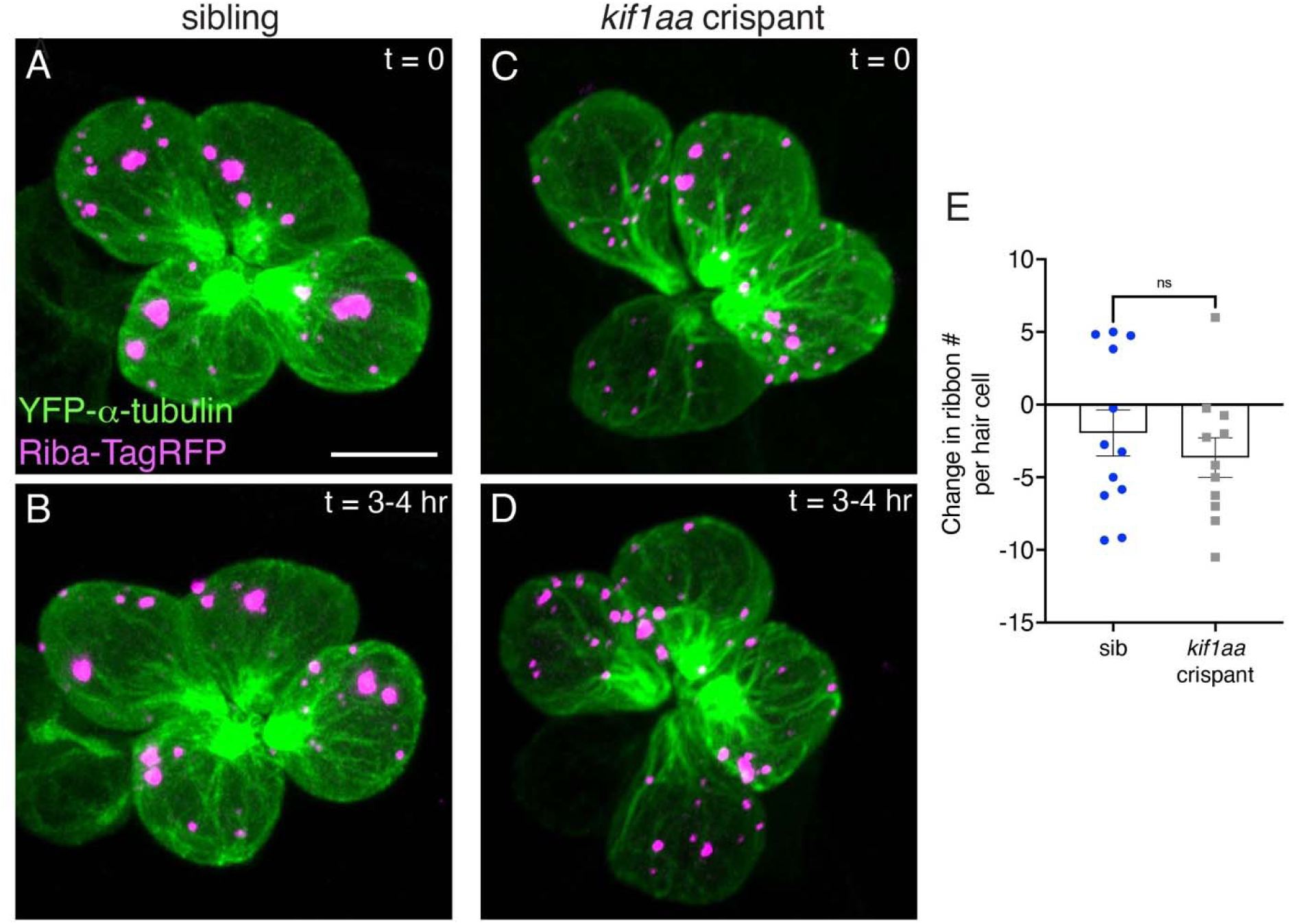
Loss of Kif1aa does not impact ribbon numbers over 3-4 hrs. **A,C)** Example images of neuromasts at 2 dpf. The microtubule network and ribbons are marked with YFP-tubulin and Riba-TagRFP respectively. Neuromasts were imaged immediately (**A**, sibling control; **C**, *kif1aa* F0 crispants) and after 3-4 hrs (**B**, sibling control; **D**, *kif1aa* F0 crispants). **E**) Quantification revealed that after 3-4 hrs the number of Riba-TagRFP puncta per hair cell was the same in control and *kif1aa* F0 crispants (n = 12 and 11 neuromasts for control and *kif1aa* F0 crispants, P = 0.427). An unpaired t-test was used for the comparison in **E**. Scale bar in **A** = 5 µm.

**Figure 7-S1.**
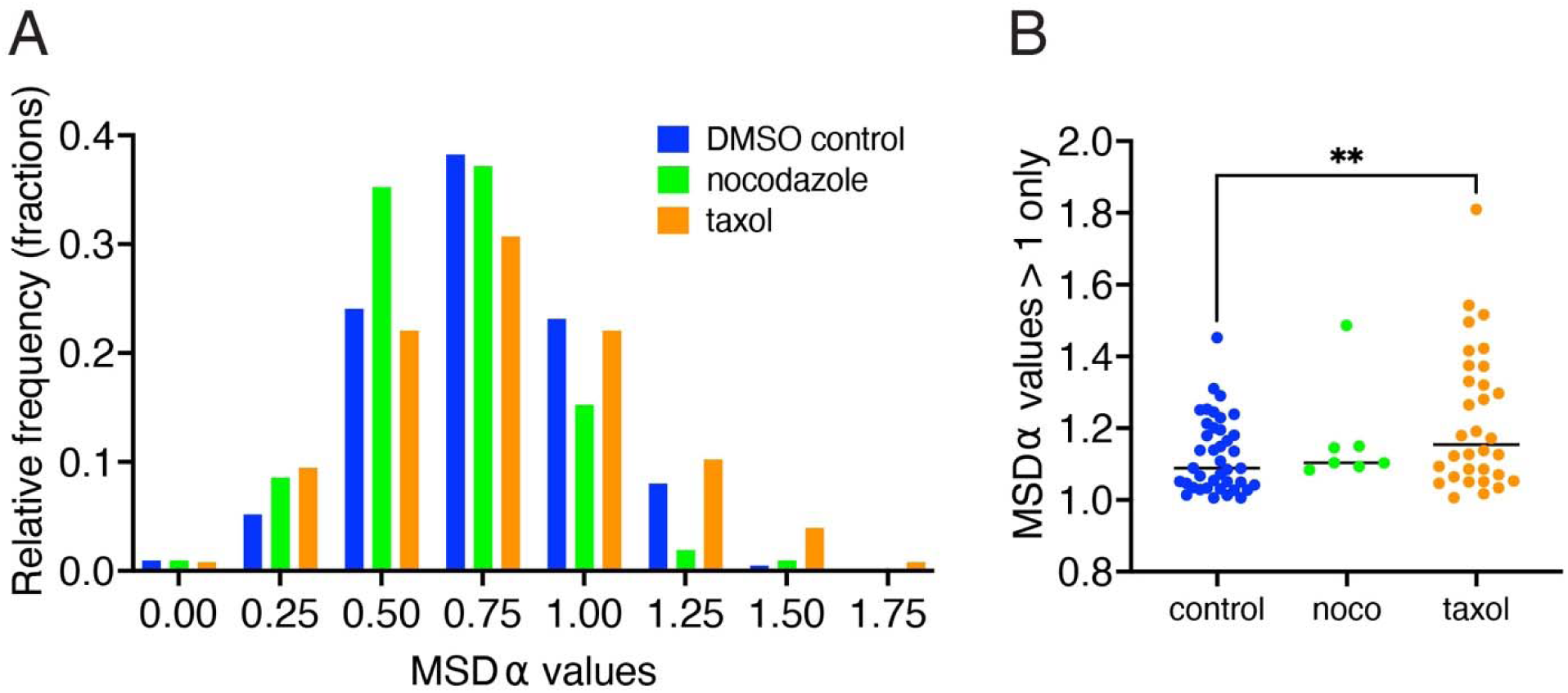
A more stable microtubule network results in directional ribbon tracks with higher MSD α values. **A** Distributions of individual ribbon track MSD α values are shown for control, nocodazole-(500 nM) and taxol-(25 µM) treated hair cells. Compared to control, taxol treatment shifts the distribution towards higher α values, while nocodazole shifts the distribution towards lower α values (total number of tracks analyzed: control (239), nocodazole (113), and taxol (140)). **B)** When examining only α > 1 values (indicative of directional motion), there are significantly higher α values in taxol-treated samples compared to control (P = 0.013). A one-way ANOVA was used for the comparison in **B**.

**Figure 8-S1.**
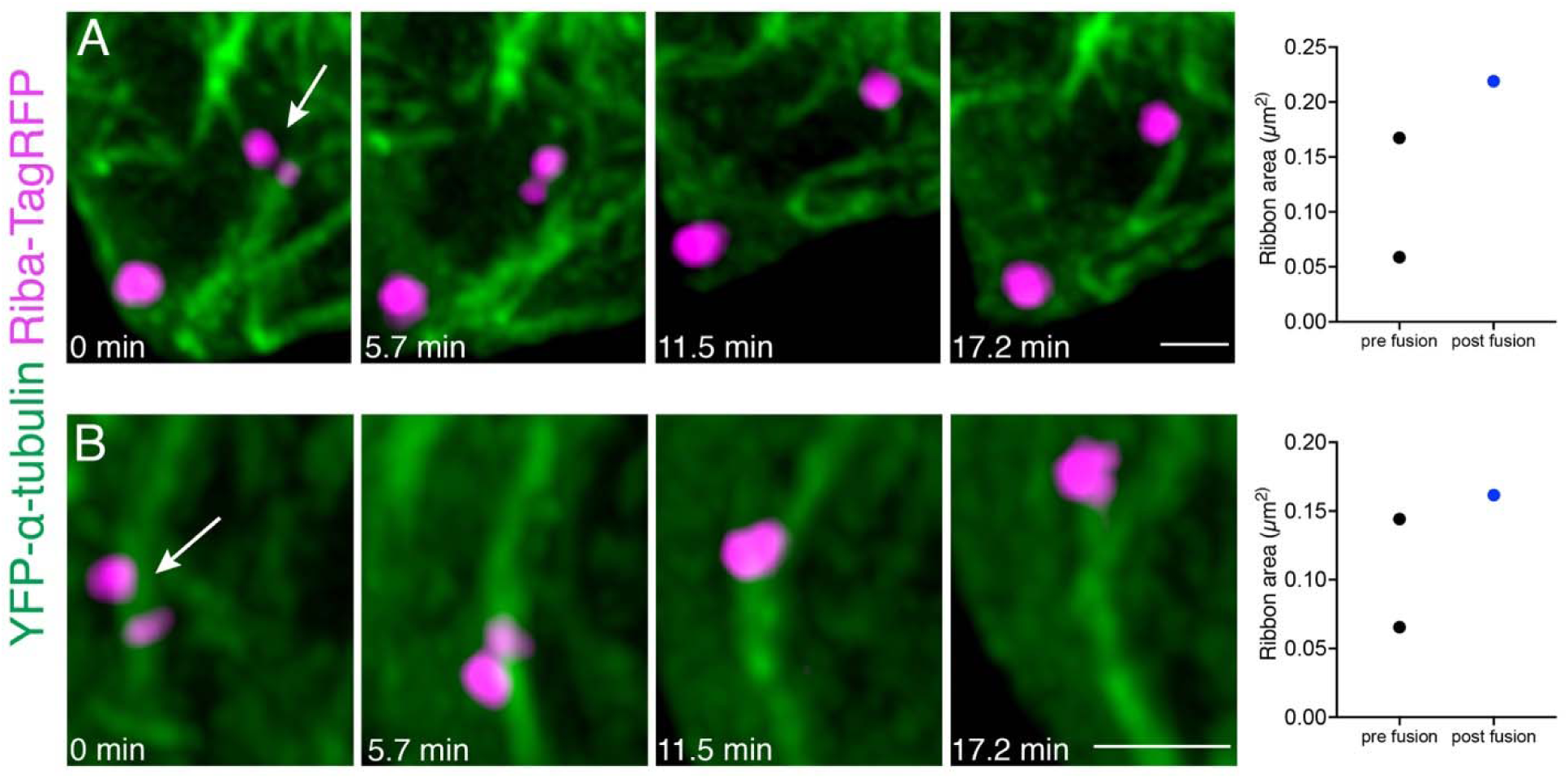
Ribbon precursor fuse on or near microtubules. **A-B)** Two examples of ribbon precursor fusion on or near microtubules. Ribbons were tracked over 13 frames that were acquired every 5.7 min. Shown in **A** are frames 3-6. Shown in **B** are frames 4-7. The arrows indicate ribbons of interest. Plotted on the right of each example is the change in area before (pre fusion) the 2 precursors fuse into one larger precursor (post fusion). These examples correspond to Movies S10 and S11. Scale bars in **A-B** = 1 µm.

## Movie Legends

**Movie S1 – EB3-GFP dynamics in developing lateral-line hair cells**

Timelapse of the example of a pLL neuromast expressing EB3-GFP from Figure 2. The growing or plus ends of microtubules can be visualized by capturing EB3-GFP dynamics. The timelapse was acquired on a LSM 780 confocal microscope every 7 s for 22 min. Maximum-intensity projection of the original z-stack is shown on the left side, played at 5 frames per second. The right side shows the same movie as the left side except EB3-GFP tracks are color-coded and followed over time using the FIJI plugin TrackMate. Scale bar = 5 µm

**Movie S2 – Tracking precursors and ribbons in 3D using Imaris**

Timelapse movie of a pLL neuromast at 2 dpf. A partial cell volume was captured at 50 s intervals for ∼30 min with a Zeiss LSM 780 with Airyscan. Microtubules are marked with YFP-tubulin (green), ribbons and precursors are marked with Riba-TagRFP (magenta). Tracks captured in Imaris are color coded with time. This example is the same as Figure 3A-B. Maximum-intensity projections of the original z-stacks are shown, played at 5 frames per second. Scale bar = 2 µm.

**Movie S3 – Tracking precursors and ribbons in 3D using Imaris**

Timelapse movie of a pLL neuromast at 2 dpf. A partial cell volume was captured at 60 s intervals for ∼30 min with a Zeiss LSM 780 with Airyscan. Microtubules are marked with YFP-tubulin (green), ribbons and precursors are marked with Riba-TagRFP (magenta). Tracks captured in Imaris are shown in yellow. Maximum-intensity projections of the original z-stacks are shown, played at 5 frames per second. Scale bar = 2 µm.

**Movie S4 – Directional motion to cell base and stationary precursors on microtubule**

Timelapse movie of a pLL neuromast at 2 dpf. A partial cell volume was captured at 20 s intervals for 9 min with a Zeiss LSM 980 with Airyscan 2. Microtubules are marked with YFP-tubulin (green), ribbons and precursors are marked with Riba-TagRFP (magenta). Maximum-intensity projections of the original z-stacks are shown played at 5 frames per second. This example is the same as Figure 3D-E. On the left side is a timelapse of the hair cell showing a ribbon precursor moving along a microtubule toward the cell base. In addition, a stationary ribbon precursor is also shown. On the right is a higher magnification view of these two ribbon precursors. The circles indicate spots identified using TrackMate, along with the track of the moving ribbon shown in yellow. Scale bar = 2 µm.

**Movie S5 – Directional motion of precursor along a microtubule to the cell base**

Timelapse movie of a pLL neuromast at 2 dpf. A partial cell volume was captured at 3 s intervals for 5 min with a Zeiss LSM 980 with Airyscan 2. Microtubules are marked with YFP-tubulin (green), ribbons and precursors are marked with Riba-TagRFP (magenta). Maximum-intensity projections of the original z-stacks are shown, played at 5 frames per second. On the left side is a timelapse of the hair cell showing a ribbon precursor moving along a microtubule toward the cell base. On the right is a higher magnification view of these two ribbon precursors. TrackMate was used to generate the track of the precursor along the microtubule in yellow. This example is the same as Figure 3-S1A. Scale bar = 2 µm.

**Movie S6 – Directional motion of precursor along microtubule to cell apex**

Timelapse movie of a pLL neuromast at 2 dpf. A partial cell volume was captured at 50 s intervals for 18 min with a Zeiss LSM 780 with Airyscan. Microtubules are marked with YFP-tubulin (green), ribbons and precursors are marked with Riba-TagRFP (magenta). Maximum-intensity projections of the original z-stacks are shown, played at 5 frames per second. On the left side is a timelapse of the hair cell showing a ribbon precursor moving along a microtubule toward the cell apex. On the right is a higher magnification view of this precursor, with the tracks of the ribbon precursor during the timelapse. The circle indicates a spot identified using TrackMate, along with the track of the moving ribbon shown in yellow. This example is the same as Figure 3-S1B. Scale bar = 2 µm.

**Movie S7 – Precursor switching between microtubules**

Timelapse movie of a pLL neuromast at 2 dpf. A partial cell volume was captured at ∼100 s intervals for 40 min with a Zeiss LSM 980 with Airyscan. Microtubules are marked with YFP-tubulin (green), ribbons and precursors are marked with Riba-TagRFP (magenta). Maximum-intensity projections of the original z-stacks are shown, played at 5 frames per second. On the left side is a timelapse of the hair cell showing a ribbon precursor moving along a microtubule toward the hair cell apex; the precursor then moves to another microtubule. On the right is a higher magnification view of the ribbon precursor during the timelapse. The precursor movement is quite fast and at this interval can be observed twice in a single z-stack during movement between microtubules at timepoints 2,4, and 15. This example is the same as Figure 3-S1C. Scale bar = 2 µm.

**Movie S8 – Microtubule dynamics in hair cells change upon treatment with nocodazole and taxol**

Timelapse movies of pLL neuromasts at 2 dpf captured after treatment with 0.1 % DMSO (control, left side), 25 µM taxol (Taxol, middle), or 250 nM nocodazole (Noc, right side) for 30 min. Microtubules are marked with YFP-tubulin (gray) and ribbons are marked with Riba-TagRFP (magenta). The timelapses of partial cell volumes were acquired with a Zeiss LSM 780 with Airyscan every 50-80 s for 30 min. Maximum-intensity projections of the original z-stacks are shown, played at 5 frames per second. The taxol-treated neuromast has more stabilized microtubules and very little cytoplasmic tubulin (depolymerized tubulin), indicating that the microtubules are more stable than the control. In the nocodazole-treated neuromast, there are fewer microtubules and more diffuse cytoplasmic tubulin compared to the control. Scale bar = 2 µm.

**Movie S9 – Ribbon precursors attached to microtubules undergo fusion**

Timelapse movie of a pLL neuromast at 2 dpf. A partial cell volume was captured at 4.8 min intervals for 100 min with a Zeiss LSM 780 with Airyscan. Microtubules are marked with YFP-tubulin (green) and ribbons are marked with Riba-TagRFP (magenta). Maximum-intensity projections of the original z-stacks are shown, played at 5 frames per second. Two ribbon precursors associate with microtubules and fuse near the hair cell base. This example is the same as Figure 8A-B. Scale bar = 2 µm.

**Movie S10 – Ribbon precursors attached to microtubules undergo fusion**

Timelapse movie of a pLL neuromast at 2 dpf. A partial cell volume was captured at 5.7 min intervals for 75 min with a Zeiss LSM 780 with Airyscan. Microtubules are marked with YFP-tubulin (green) and ribbons are marked with Riba-TagRFP (magenta). Maximum-intensity projections of the original z-stacks are shown, played at 2 frames per second. This example is the same as Figure 8-S1A. Scale bar = 2 µm.

**Movie 11 – Ribbon precursors attached to microtubules undergo fusion**

Timelapse movie of a pLL neuromast at 2 dpf. A partial cell volume was captured at 5.7 min intervals for 75 min with a Zeiss LSM 780 with Airyscan. Microtubules are marked with YFP-tubulin (green) and ribbons are marked with Riba-TagRFP (magenta). Maximum-intensity projections of the original z-stacks are shown, played at 2 frames per second. This example is the same as Figure 8-S1B. Scale bar = 2 µm.

## References

Adamíková L, Straube A, Schulz I, Steinberg G. 2004. Calcium signaling is involved in dynein-dependent microtubule organization. Mol Biol Cell 15:1969–1980. doi:10.1091/mbc.E03-09-0675

Ahmari SE, Buchanan J, Smith SJ. 2000. Assembly of presynaptic active zones from cytoplasmic transport packets. Nat Neurosci 3:445–451. doi:10.1038/74814

Becker L, Schnee ME, Niwa M, Sun W, Maxeiner S, Talaei S, Kachar B, Rutherford MA, Ricci AJ. 2018. The presynaptic ribbon maintains vesicle populations at the hair cell afferent fiber synapse. eLife 7:e30241. doi:10.7554/eLife.30241

Bellotti A, Murphy J, Lin L, Petralia R, Wang Y-X, Hoffman D, O’Leary T. 2021. Paradoxical relationships between active transport and global protein distributions in neurons. Biophys J 120:2085–2101. doi:10.1016/j.bpj.2021.02.048

Brandt A, Striessnig J, Moser T. 2003. CaV1.3 channels are essential for development and presynaptic activity of cochlear inner hair cells. J Neurosci 23:10832–10840.

Bury LAD, Sabo SL. 2011. Coordinated trafficking of synaptic vesicle and active zone proteins prior to synapse formation. Neural Develop 6:24. doi:10.1186/1749-8104-6-24

Carrington B, Varshney GK, Burgess SM, Sood R. 2015. CRISPR-STAT: an easy and reliable PCR-based method to evaluate target-specific sgRNA activity. Nucleic Acids Res 43:e157. doi:10.1093/nar/gkv802

Ceriani F, Hendry A, Jeng J-Y, Johnson SL, Stephani F, Olt J, Holley MC, Mammano F, Engel J, Kros CJ, Simmons DD, Marcotti W. 2019. Coordinated calcium signalling in cochlear sensory and non-sensory cells refines afferent innervation of outer hair cells. EMBO J 38:e99839. doi:10.15252/embj.201899839

Chang B, Heckenlively JR, Bayley PR, Brecha NC, Davisson MT, Hawes NL, Hirano AA, Hurd RE, Ikeda A, Johnson BA, Mccall MA, Morgans CW, Nusinowitz S, Peachey NS, Rice DS, Vessey KA, Gregg RG. 2006. The nob2 mouse, a null mutation in Cacna1f: Anatomical and functional abnormalities in the outer retina and their consequences on ganglion cell visual responses. Vis Neurosci 23:11–24. doi:10.1017/S095252380623102X

Chernov KG, Barbet A, Hamon L, Ovchinnikov LP, Curmi PA, Pastré D. 2009. Role of microtubules in stress granule assembly. J Biol Chem 284:36569–36580. doi:10.1074/jbc.M109.042879

Cochard A, Safieddine A, Combe P, Benassy M-N, Weil D, Gueroui Z. 2023. Condensate functionalization with microtubule motors directs their nucleation in space and allows manipulating RNA localization. EMBO J n/a:e114106. doi:10.15252/embj.2023114106

Corradi E, Dalla Costa I, Gavoci A, Iyer A, Roccuzzo M, Otto TA, Oliani E, Bridi S, Strohbuecker S, Santos-Rodriguez G, Valdembri D, Serini G, Abreu-Goodger C, Baudet M. 2020. Axonal precursor miRNAs hitchhike on endosomes and locally regulate the development of neural circuits. EMBO J 39:e102513. doi:10.15252/embj.2019102513

David S, Pinter K, Nguyen K-K, Lee DS, Lei Z, Sokolova Y, Sheets L, Kindt KS. 2024. Kif1a and intact microtubules maintain synaptic-vesicle populations at ribbon synapses in zebrafish hair cells. J Physiol. doi:10.1113/JP286263

Dick O, Tom Dieck S, Altrock WD, Ammermüller J, Weiler R, Garner CC, Gundelfinger ED, Brandstätter JH. 2003. The presynaptic active zone protein bassoon is essential for photoreceptor ribbon synapse formation in the retina. Neuron 37:775–786. doi:10.1016/s0896-6273(03)00086-2

Dow E, Siletti K, Hudspeth AJ. 2015. Cellular projections from sensory hair cells form polarity-specific scaffolds during synaptogenesis. Genes Dev 29:1087–1094. doi:10.1101/gad.259838.115

Façanha ALO, Appelgren H, Tabish M, Okorokov L, Ekwall K. 2002. The endoplasmic reticulum cation P-type ATPase Cta4p is required for control of cell shape and microtubule dynamics. J Cell Biol 157:1029–1039. doi:10.1083/jcb.200111012

Farías GG, Cuitino L, Guo X, Ren X, Jarnik M, Mattera R, Bonifacino JS. 2012. Signal-mediated, AP-1/clathrin-dependent sorting of transmembrane receptors to the somatodendritic domain of hippocampal neurons. Neuron 75:810–823. doi:10.1016/j.neuron.2012.07.007

Fejtova A, Davydova D, Bischof F, Lazarevic V, Altrock WD, Romorini S, Schöne C, Zuschratter W, Kreutz MR, Garner CC, Ziv NE, Gundelfinger ED. 2009. Dynein light chain regulates axonal trafficking and synaptic levels of Bassoon. J Cell Biol 185:341–355. doi:10.1083/jcb.200807155

Frank T, Rutherford MA, Strenzke N, Neef A, Pangršič T, Khimich D, Fejtova A, Gundelfinger ED, Liberman MC, Harke B, Bryan KE, Lee A, Egner A, Riedel D, Moser T. 2010. Bassoon and the synaptic ribbon organize Ca^2^+ channels and vesicles to add release sites and promote refilling. Neuron 68:724–738. doi:10.1016/j.neuron.2010.10.027

Frederick CE, Zenisek D. 2023. Ribbon synapses and retinal disease: review. Int J Mol Sci 24:5090. doi:10.3390/ijms24065090

Freeman W. 1928. The function of the lateral line organs. Science 68:205–205. doi:10.1126/science.68.1757.205

Fu M, Holzbaur ELF. 2014. Integrated regulation of motor-driven organelle transport by scaffolding proteins. Trends Cell Biol 24:564–574. doi:10.1016/j.tcb.2014.05.002

Gerdes HH, Rosa P, Phillips E, Baeuerle PA, Frank R, Argos P, Huttner WB. 1989. The primary structure of human secretogranin II, a widespread tyrosine-sulfated secretory granule protein that exhibits low pH- and calcium-induced aggregation. J Biol Chem 264:12009– 12015. doi:10.1016/S0021-9258(18)80167-3

Graydon CW, Manor U, Kindt KS. 2017. In vivo ribbon mobility and turnover of Ribeye at zebrafish hair cell synapses. Sci Rep 7:7467. doi:10.1038/s41598-017-07940-z

Guillet M, Sendin G, Bourien J, Puel J-L, Nouvian R. 2016. Actin filaments regulate exocytosis at the hair cell ribbon synapse. J Neurosci 36:649–654. doi:10.1523/JNEUROSCI.3379-15.2016

Gundelfinger ED, Reissner C, Garner CC. 2016. Role of bassoon and piccolo in assembly and molecular organization of the active zone. Front Synaptic Neurosci 7:19. doi:10.3389/fnsyn.2015.00019

Gupta RS. 1985. Species-specific differences in toxicity of antimitotic agents toward cultured mammalian cells2. JNCI J Natl Cancer Inst 74:159–164. doi:10.1093/jnci/74.1.159

Hoshijima K, Jurynec MJ, Klatt Shaw D, Jacobi AM, Behlke MA, Grunwald DJ. 2019. Highly efficient crispr-cas9-based methods for generating deletion mutations and f0 embryos that lack gene function in zebrafish. Dev Cell 51:645–657.e4. doi:10.1016/j.devcel.2019.10.004

Hummel JJA, Hoogenraad CC. 2021. Specific KIF1A-adaptor interactions control selective cargo recognition. J Cell Biol 220:e202105011. doi:10.1083/jcb.202105011

Janke C, Magiera MM. 2020. The tubulin code and its role in controlling microtubule properties and functions. Nat Rev Mol Cell Biol 21:307–326. doi:10.1038/s41580-020-0214-3

Jean P, Lopez de la Morena D, Michanski S, Jaime Tobón LM, Chakrabarti R, Picher MM, Neef J, Jung S, Gültas M, Maxeiner S, Neef A, Wichmann C, Strenzke N, Grabner C, Moser T. 2018. The synaptic ribbon is critical for sound encoding at high rates and with temporal precision. eLife 7:e29275. doi:10.7554/eLife.29275

Jing Z, Rutherford MA, Takago H, Frank T, Fejtova A, Khimich D, Moser T, Strenzke N. 2013. Disruption of the presynaptic cytomatrix protein bassoon degrades ribbon anchorage, multiquantal release, and sound encoding at the hair cell afferent synapse. J Neurosci 33:4456–67. doi:10.1523/JNEUROSCI.3491-12.2013

Kawano D, Pinter K, Chlebowski M, Petralia RS, Wang Y-X, Nechiporuk AV, Drerup CM. 2022. NudC regulated Lis1 stability is essential for the maintenance of dynamic microtubule ends in axon terminals. iScience 25:105072. doi:10.1016/j.isci.2022.105072

Khimich D, Nouvian R, Pujol R, Tom Dieck S, Egner A, Gundelfinger ED, Moser T. 2005. Hair cell synaptic ribbons are essential for synchronous auditory signalling. Nature 434:889–894. doi:10.1038/nature03418

Kindt KS, Finch G, Nicolson T. 2012. Kinocilia mediate mechanosensitivity in developing zebrafish hair cells. Dev Cell 23:329–341. doi:10.1016/j.devcel.2012.05.022

Kujawa SG, Liberman MC. 2015. Synaptopathy in the noise-exposed and aging cochlea: primary neural degeneration in acquired sensorineural hearing loss. Hear Res 330:191–199. doi:10.1016/j.heares.2015.02.009

Kwan KM, Fujimoto E, Grabher C, Mangum BD, Hardy ME, Campbell DS, Parant JM, Yost HJ, Kanki JP, Chien C-B. 2007. The Tol2kit: a multisite gateway-based construction kit for Tol2 transposon transgenesis constructs. Dev Dyn Off Publ Am Assoc Anat 236:3088– 3099. doi:10.1002/dvdy.21343

Laisne M-C, Michallet S, Lafanechère L. 2021. Characterization of microtubule destabilizing drugs: a quantitative cell-based assay that bridges the gap between tubulin based- and cytotoxicity assays. Cancers 13:5226. doi:10.3390/cancers13205226

Leighton AH, Lohmann C. 2016. The wiring of developing sensory circuits-from patterned spontaneous activity to synaptic plasticity mechanisms. Front Neural Circuits 10:71. doi:10.3389/fncir.2016.00071

Lepelletier L, de Monvel JB, Buisson J, Desdouets C, Petit C. 2013. Auditory hair cell centrioles undergo confined brownian motion throughout the developmental migration of the kinocilium. Biophys J 105:48–58. doi:10.1016/j.bpj.2013.05.009

Luo Y-Y, Wu J-J, Li Y-M. 2021. Regulation of liquid–liquid phase separation with focus on post-translational modifications. Chem Commun 57:13275–13287. doi:10.1039/D1CC05266G

Lush ME, Diaz DC, Koenecke N, Baek S, Boldt H, St Peter MK, Gaitan-Escudero T, Romero-Carvajal A, Busch-Nentwich EM, Perera AG, Hall KE, Peak A, Haug JS, Piotrowski T. 2019. scRNA-Seq reveals distinct stem cell populations that drive hair cell regeneration after loss of Fgf and Notch signaling. eLife 8:e44431. doi:10.7554/eLife.44431

Lv C, Stewart WJ, Akanyeti O, Frederick C, Zhu J, Santos-Sacchi J, Sheets L, Liao JC, Zenisek D. 2016. Synaptic ribbons require ribeye for electron density, proper synaptic localization, and recruitment of calcium channels. Cell Rep 15:2784–2795. doi:10.1016/j.celrep.2016.05.045

Maas C, Torres VI, Altrock WD, Leal-Ortiz S, Wagh D, Terry-Lorenzo RT, Fejtova A, Gundelfinger ED, Ziv NE, Garner CC. 2012. Formation of Golgi-derived active zone precursor vesicles. J Neurosci 32:11095–11108. doi:10.1523/JNEUROSCI.0195-12.2012

Magupalli VG, Schwarz K, Alpadi K, Natarajan S, Seigel GM, Schmitz F. 2008. Multiple RIBEYE-RIBEYE interactions create a dynamic scaffold for the formation of synaptic ribbons. J Neurosci 28:7954–7967. doi:10.1523/JNEUROSCI.1964-08.2008

Maxeiner S, Luo F, Tan A, Schmitz F, Südhof TC. 2016. How to make a synaptic ribbon: RIBEYE deletion abolishes ribbons in retinal synapses and disrupts neurotransmitter release. EMBO J 35:1098–1114. doi:10.15252/embj.201592701

Merriam EB, Millette M, Lumbard DC, Saengsawang W, Fothergill T, Hu X, Ferhat L, Dent EW. 2013. Synaptic regulation of microtubule dynamics in dendritic spines by calcium, F-actin, and drebrin. J Neurosci 33:16471–16482. doi:10.1523/JNEUROSCI.0661-13.2013

Michanski S, Kapoor R, Steyer AM, Möbius W, Früholz I, Ackermann F, Gültas M, Garner CC, Hamra FK, Neef J, Strenzke N, Moser T, Wichmann C. 2023. Piccolino is required for ribbon architecture at cochlear inner hair cell synapses and for hearing. EMBO Rep n/a:e56702. doi:10.15252/embr.202256702

Michanski S, Smaluch K, Steyer AM, Chakrabarti R, Setz C, Oestreicher D, Fischer C, Möbius W, Moser T, Vogl C, Wichmann C. 2019. Mapping developmental maturation of inner hair cell ribbon synapses in the apical mouse cochlea. Proc Natl Acad Sci 116:6415–6424. doi:10.1073/pnas.1812029116

Niwa S, Tanaka Y, Hirokawa N. 2008. KIF1Bβ- and KIF1A-mediated axonal transport of presynaptic regulator Rab3 occurs in a GTP-dependent manner through DENN/MADD. Nat Cell Biol 10:1269–1279. doi:10.1038/ncb1785

Obholzer N, Wolfson S, Trapani JG, Mo W, Nechiporuk A, Busch-Nentwich E, Seiler C, Sidi S, Söllner C, Duncan RN, Boehland A, Nicolson T. 2008. Vesicular glutamate transporter 3 is required for synaptic transmission in zebrafish hair cells. J Neurosci 28:2110–2118. doi:10.1523/JNEUROSCI.5230-07.2008

Ohta S, Ji YR, Martin D, Wu DK. 2020. Emx2 regulates hair cell rearrangement but not positional identity within neuromasts. eLife. doi:10.7554/eLife.60432

Okada Y, Yamazaki H, Sekine-Aizawa Y, Hirokawa N. 1995. The neuron-specific kinesin superfamily protein KIF1A is a unique monomeric motor for anterograde axonal transport of synaptic vesicle precursors. Cell 81:769–780. doi:10.1016/0092-8674(95)90538-3

Oliver D, Ramachandran S, Philbrook A, Lambert CM, Nguyen KCQ, Hall DH, Francis MM. 2022. Kinesin-3 mediated axonal delivery of presynaptic neurexin stabilizes dendritic spines and postsynaptic components. PLoS Genet 18:e1010016. doi:10.1371/journal.pgen.1010016

Pack-Chung E, Kurshan PT, Dickman DK, Schwarz TL. 2007. A Drosophila kinesin required for synaptic bouton formation and synaptic vesicle transport. Nat Neurosci 10:980–989. doi:10.1038/nn1936

Parslow A, Cardona A, Bryson-Richardson RJ. 2014. Sample drift correction following 4D confocal time-lapse imaging. J Vis Exp JoVE 51086. doi:10.3791/51086

Regus-Leidig H, Fuchs M, Löhner M, Leist SR, Leal-Ortiz S, Chiodo VA, Hauswirth WW, Garner CC, Brandstätter JH. 2014. In vivo knockdown of Piccolino disrupts presynaptic ribbon morphology in mouse photoreceptor synapses. Front Cell Neurosci 8.

Regus-Leidig H, Tom Dieck S, Specht D, Meyer L, Brandstätter JH. 2009. Early steps in the assembly of photoreceptor ribbon synapses in the mouse retina: The involvement of precursor spheres. J Comp Neurol 512:814–824. doi:10.1002/cne.21915

Roux I, Hosie S, Johnson SL, Bahloul A, Cayet N, Nouaille S, Kros CJ, Petit C, Safieddine S. 2009. Myosin VI is required for the proper maturation and function of inner hair cell ribbon synapses. Hum Mol Genet 18:4615–4628. doi:10.1093/hmg/ddp429

Ruel J, Emery S, Nouvian R, Bersot T, Amilhon B, Van Rybroek JM, Rebillard G, Lenoir M, Eybalin M, Delprat B, Sivakumaran TA, Giros B, El Mestikawy S, Moser T, Smith RJH, Lesperance MM, Puel J-L. 2008. Impairment of SLC17A8 encoding vesicular glutamate transporter-3, VGLUT3, underlies nonsyndromic deafness DFNA25 and inner hair cell dysfunction in null mice. Am J Hum Genet 83:278–292. doi:10.1016/j.ajhg.2008.07.008

Sampo B, Kaech S, Kunz S, Banker G. 2003. Two distinct mechanisms target membrane proteins to the axonal surface. Neuron 37:611–624. doi:10.1016/s0896-6273(03)00058-8

Schmitz F. 2009. The making of synaptic ribbons: how they are built and what they do. Neurosci Rev J Bringing Neurobiol Neurol Psychiatry 15:611–624. doi:10.1177/1073858409340253

Schmitz F, Königstorfer A, Südhof TC. 2000. RIBEYE, a component of synaptic ribbons: a protein’s journey through evolution provides insight into synaptic ribbon function. Neuron 28:857–872. doi:10.1016/S0896-6273(00)00159-8

Schrøder JM, Larsen J, Komarova Y, Akhmanova A, Thorsteinsson RI, Grigoriev I, Manguso R, Christensen ST, Pedersen SF, Geimer S, Pedersen LB. 2011. EB1 and EB3 promote cilia biogenesis by several centrosome-related mechanisms. J Cell Sci 124:2539–2551. doi:10.1242/jcs.085852

Shapira M, Zhai RG, Dresbach T, Bresler T, Torres VI, Gundelfinger ED, Ziv NE, Garner CC. 2003. Unitary assembly of presynaptic active zones from Piccolo-Bassoon transport vesicles. Neuron 38:237–252. doi:10.1016/s0896-6273(03)00207-1

Sheets L, Holmgren M, Kindt KS. 2021. How zebrafish can drive the future of genetic-based hearing and balance research. JARO J Assoc Res Otolaryngol 22:215–235. doi:10.1007/s10162-021-00798-z

Sheets L, Kindt KS, Nicolson T. 2012. Presynaptic CaV1.3 channels regulate synaptic ribbon size and are required for synaptic maintenance in sensory hair cells. J Neurosci 32:17273– 17286. doi:10.1523/JNEUROSCI.3005-12.2012

Sheets L, Trapani JG, Mo W, Obholzer N, Nicolson T. 2011. Ribeye is required for presynaptic Ca(V)1.3a channel localization and afferent innervation of sensory hair cells. Dev Camb Engl 138:1309–1319. doi:10.1242/dev.059451

Sikora G, Teuerle M, Wyłomańska A, Grebenkov D. 2017. Statistical properties of the anomalous scaling exponent estimator based on time-averaged mean-square displacement. Phys Rev E 96:022132. doi:10.1103/PhysRevE.96.022132

Sobkowicz HM, Rose JE, Scott GE, Slapnick SM. 1982. Ribbon synapses in the developing intact and cultured organ of Corti in the mouse. J Neurosci 2:942–957. doi:10.1523/JNEUROSCI.02-07-00942.1982

Sobkowicz HM, Rose JE, Scott GL, Levenick CV. 1986. Distribution of synaptic ribbons in the developing organ of Corti. J Neurocytol 15:693–714. doi:10.1007/BF01625188

Stepanova T, Slemmer J, Hoogenraad CC, Lansbergen G, Dortland B, Zeeuw CID, Grosveld F, Cappellen G van, Akhmanova A, Galjart N. 2003. Visualization of microtubule growth in cultured neurons via the use of eb3-gfp (end-binding protein 3-green fluorescent protein). J Neurosci 23:2655–2664. doi:10.1523/JNEUROSCI.23-07-02655.2003

Suli A, Watson GM, Rubel EW, Raible DW. 2012. Rheotaxis in larval zebrafish is mediated by lateral line mechanosensory hair cells. PLoS ONE 7. doi:10.1371/journal.pone.0029727

Sur A, Wang Y, Capar P, Margolin G, Prochaska MK, Farrell JA. 2023. Single-cell analysis of shared signatures and transcriptional diversity during zebrafish development. Dev Cell 58:3028–3047.e12. doi:10.1016/j.devcel.2023.11.001

Sweeney HL, Holzbaur ELF. 2018. Motor Proteins. Cold Spring Harb Perspect Biol 10:a021931. doi:10.1101/cshperspect.a021931

Tarantino N, Tinevez J-Y, Crowell EF, Boisson B, Henriques R, Mhlanga M, Agou F, Israël A, Laplantine E. 2014. TNF and IL-1 exhibit distinct ubiquitin requirements for inducing NEMO–IKK supramolecular structures. J Cell Biol 204:231–245. doi:10.1083/jcb.201307172

Tinevez J-Y, Perry N, Schindelin J, Hoopes GM, Reynolds GD, Laplantine E, Bednarek SY, Shorte SL, Eliceiri KW. 2017. TrackMate: An open and extensible platform for single-particle tracking. *Methods*, Image Processing for Biologists 115:80–90. doi:10.1016/j.ymeth.2016.09.016

Trapani JG, Obholzer N, Mo W, Brockerhoff SE, Nicolson T. 2009. Synaptojanin1 is required for temporal fidelity of synaptic transmission in hair cells. PLOS Genet 5:e1000480. doi:10.1371/journal.pgen.1000480

Tritsch NX, Yi E, Gale JE, Glowatzki E, Bergles DE. 2007. The origin of spontaneous activity in the developing auditory system. Nature 450:50–55. doi:10.1038/nature06233

Uthaiah RC, Hudspeth AJ. 2010. Molecular anatomy of the hair cell’s ribbon synapse. J Neurosci 30:12387–12399. doi:10.1523/JNEUROSCI.1014-10.2010

Varshney GK, Carrington B, Pei W, Bishop K, Chen Z, Fan C, Xu L, Jones M, LaFave MC, Ledin J, Sood R, Burgess SM. 2016. A high-throughput functional genomics workflow based on CRISPR/Cas9-mediated targeted mutagenesis in zebrafish. Nat Protoc 11:2357–2375. doi:10.1038/nprot.2016.141

Vleugel M, Kok M, Dogterom M. 2016. Understanding force-generating microtubule systems through in vitro reconstitution. Cell Adhes Migr 10:475–494. doi:10.1080/19336918.2016.1241923

Voorn RA, Sternbach M, Jarysta A, Rankovic V, Tarchini B, Wolf F, Vogl C. 2024. Slow kinesin-dependent microtubular transport facilitates ribbon synapse assembly in developing cochlear inner hair cells. eLife 13. doi:10.7554/eLife.98145.1

Wan G, Ji L, Schrepfer T, Gong S, Wang G-P, Corfas G. 2019. Synaptopathy as a mechanism for age-related vestibular dysfunction in mice. Front Aging Neurosci 11:156. doi:10.3389/fnagi.2019.00156

Wang B, Zhang Lei, Dai T, Qin Z, Lu H, Zhang Long, Zhou F. 2021. Liquid–liquid phase separation in human health and diseases. Signal Transduct Target Ther 6:1–16. doi:10.1038/s41392-021-00678-1

Wang Z, Lou J, Zhang H. 2022. Essence determines phenomenon: Assaying the material properties of biological condensates. J Biol Chem 298. doi:10.1016/j.jbc.2022.101782

Wiegand T, Hyman AA. 2020. Drops and fibers — how biomolecular condensates and cytoskeletal filaments influence each other. Emerg Top Life Sci 4:247–261. doi:10.1042/ETLS20190174

Wong HC, Zhang Q, Beirl AJ, Petralia RS, Wang Y-X, Kindt K. 2019. Synaptic mitochondria regulate hair-cell synapse size and function. eLife 8:e48914. doi:10.7554/eLife.48914

Yamada M, Tanaka-Takiguchi Y, Hayashi M, Nishina M, Goshima G. 2017. Multiple kinesin-14 family members drive microtubule minus end–directed transport in plant cells. J Cell Biol 216:1705–1714. doi:10.1083/jcb.201610065

Yoo SH. 1995. Ph- and ca2+-induced conformational change and aggregation of chromogranin b.: comparison with chromogranin a and implication in secretory vesicle biogenesis (*). J Biol Chem 270:12578–12583. doi:10.1074/jbc.270.21.12578

Zhang Q, Kindt KS. 2022. Using light-sheet microscopy to study spontaneous activity in the developing lateral-line system. Front Cell Dev Biol 10:819612. doi:10.3389/fcell.2022.819612

Zieve GW, Turnbull D, Mullins JM, McIntosh JR. 1980. Production of large numbers of mitotic mammalian cells by use of the reversible microtubule inhibitor nocodazole. Nocodazole accumulated mitotic cells. Exp Cell Res 126:397–405. doi:10.1016/0014-4827(80)90279-7

